# Discrepancies between ChIP-seq and CUT&Tag histone mark profiles are explained by GC content and chromatin accessibility

**DOI:** 10.64898/2026.09.22.753564

**Authors:** Eduardo Modolo, Lauren Patel, Oren Ram, Itamar Simon, Eric Mendenhall, Sven Heinz, Christopher Benner, Alon Goren

**Author notes:** Correspondence to: Alon Goren –.

## Abstract

Chromatin profiling methods, such as ChIP-seq (chromatin immunoprecipitation followed by sequencing), are used to characterize the genomic localization of DNA-associated proteins. While ChIP and ChIP-seq have been used for decades, an orthogonal approach, CUT&Tag (Cleavage Under Targets and Tagmentation), is gaining popularity as an efficient and cost-effective alternative. Although previous comparative studies note discrepancies in signal-to-noise ratios and detection bias at certain genomic regions, many differences between ChIP-seq and CUT&Tag results remain largely unreconciled. Here, we systematically investigate the disagreeing signals captured by these two methods across well annotated genomic regions. We assess multiple histone mark profiles generated by different groups in two cell lines (K562 and MCF-7). Overall, our analysis indicates that compared to ChIP-seq, CUT&Tag may have limited sensitivity in low-GC environments and, as previously observed^1–3^, increased signal in hyper-accessible chromatin. Notably, within GC-poor regions of active gene bodies and Polycomb-repressed domains, CUT&Tag exhibits a loss of H3K36me3 and H3K27me3 signal, respectively, where occupancy of these histone marks is otherwise expected. Further, active promoters with discrepant H3K4me3 and H3K27ac signal between the two assays differ systematically in GC content and chromatin accessibility. Promoters differentially enriched for CUT&Tag signal relative to ChIP-seq generally show higher, broader GC-content profiles and higher DNase-seq and ATAC-seq signal, while promoters enriched for ChIP-seq signal harbor the opposite characteristics. A local bias for high GC content and/or chromatin accessibility in CUT&Tag may also explain its differing signal patterns at nucleosome-depleted regions (NDRs) compared to ChIP-seq and MNase-seq promoter profiles. Altogether, our results highlight that studies focused on profiling the intensity and structure of histone modification occupancy can be sensitive to potential biases of CUT&Tag to GC content and chromatin accessibility. These characteristics should be accounted for when selecting a chromatin profiling approach, analyzing and interpreting data as well as drawing biological conclusions.

## Introduction

Chromatin profiling methods are used to quantify and localize DNA-associated proteins across the genome. The accurate and robust implementation of these methods have been fundamental to understanding genomic regulation. ChIP-seq^4,5^ (chromatin immunoprecipitation followed by sequencing) has been used for almost 20 years as the standard method for mapping DNA-associated proteins, with the original process of ChIP established over 40 years ago^6^. An orthogonal approach for chromatin profiling, CUT&Tag (Cleavage Under Targets and Tagmentation), has been quickly rising in popularity amongst research groups, in part due to its efficient and cost-effective protocols, compatibility with low starting material, and applications in single-cell omics^7,8^

In the commonly used ChIP-seq protocol, cells are fixed with a crosslinker such as formaldehyde^9^ and the chromatin is then sonicated and solubilized. The target protein-DNA complexes are immunoprecipitated with an antibody specific to the protein of interest, and the recovered DNA fragments are then prepared as a sequencing library. In contrast, CUT&Tag is an enzyme-tethering method performed on native, or lightly crosslinked, cells or nuclei where antibodies are added *in situ* after permeabilization. CUT&Tag builds on the Tn5 transposase enzyme^10^ which is also employed in ATAC-seq (Assay for Transposase-Accessible Chromatin) a highly used method to detect hyper-accessible chromatin^11^. In CUT&Tag, following the addition of primary and secondary antibodies, a hyperactive Tn5 transposase-Protein A fusion is targeted to the antibody complex bound to the protein of interest. The Tn5 transposase is activated, and pre-loaded DNA adapters are subsequently integrated into the DNA near the binding site of the protein of interest. The tagmented DNA fragments are then used to generate sequencing libraries^7,8^.

By leveraging distinct molecular biology compared to ChIP-seq, the CUT&Tag protocol allows for increased efficiency and generally much higher signal-to-noise ratios, as well as reduced sequencing depth and minimized requirements for starting material. Further, the retention of DNA fragments within intact cells renders CUT&Tag particularly well-suited for scalable single-cell profiling and multi-omic applications^7,12^. Together, the efficiency, ease of use and adaptability to protocol extensions including Multi-CUT&Tag^13^ and Paired-Tag^14^ are key features of CUT&Tag making it a highly valuable chromatin profiling strategy.

However, while generally producing similar results, several recent studies have reported disagreements in the output data and biological conclusions drawn between ChIP-seq and CUT&Tag^1–3,15–18^. For example, Hu et al.^3^ reports pervasive open-chromatin biases across many publicly available CUT&Tag datasets resulting in spurious signals and false-positive peaks. Abbasova et al.^1^ found that CUT&Tag datasets for histone marks H3K27ac and H3K27me3, on average, only capture about half of previously established ENCODE ChIP-seq peaks. Wang et al.^2^ reports that while CUT&Tag is particularly effective at profiling the active histone mark H3K4me3, it shows reduced efficiency compared to ChIP-seq in detecting H3K27me3 signal, which the authors attribute to lack of CUT&Tag sensitivity in compact chromatin regions. However, this observation seemingly conflicts with a report from Park et al.^15^ which finds CUT&Tag is more robust than ChIP-seq for profiling H3K9me3 in condensed heterochromatin regions. These findings imply that technical biases distinguishing ChIP-seq and CUT&Tag may prevent the methods from being fully interchangeable. In light of the apparent discrepancies and inconsistent reports regarding ChIP-seq and CUT&Tag results across recent literature, here we present a comparative analysis between the two methods to identify and characterize potential sources of differential signals.

By anchoring our analysis to well-annotated functional genomic regions (active and inactive promoters, active gene bodies, and Polycomb-repressed domains), we leverage differential enrichment analysis (DESeq2^19^) as well as relevant genomic metrics and biological readouts to identify and characterize loci where ChIP-seq and CUT&Tag profiles disagree. Importantly, we employ orthogonal and complementary reference datasets (e.g., precision nuclear run-on and sequencing (PRO-seq)^20^) to allow us to infer, in each case, which of the two profiling methods is more likely reporting the correct underlying biology. Our analysis focuses on functional regions expected to harbor well-studied histone modifications: H3K4me3, H3K27ac, H3K27me3 and H3K36me3. These histone marks encompass a range of chromatin environments and profile structures, including active or repressed transcription and narrow or broad signal distributions.

Our systematic comparison reveals that differential signals between ChIP-seq and CUT&Tag are associated with variations in genomic GC content and chromatin accessibility (as measured by DNase I hypersensitive site sequencing (DNase-seq^21^ and ATAC-seq). Specifically, we observe lower GC content and reduced chromatin accessibility signal in regions displaying limited CUT&Tag sensitivity compared to ChIP-seq, both for H3K36me3 (in active gene bodies) and for H3K27me3 (in Polycomb-repressed domains). These CUT&Tag-depleted loci retain signals such as PRO-seq and EZH2 ChIP-seq, respectively, suggesting that the observed loss of CUT&Tag signal may reflect a technical false negative at these regions. Further, we observe that active promoters with differentially-enriched H3K4me3 and H3K27ac ChIP-seq signal relative to CUT&Tag, exhibit lower, narrower GC content profiles and reduced DNase-seq and ATAC-seq signals around the transcription start site (TSS). On the other hand, promoters enriched in CUT&Tag signal relative to ChIP-seq show the opposite characteristics. Additionally, while the nucleosome-depleted region (NDR) structure at active promoters^22^ is clearly detected in MNase-seq and ChIP-seq, in a similar manner to observations made by the transposase-based chromatin profiling approach (ChIL–seq)^23^, we note that CUT&Tag profiles have a distinct signal patterns at these promoter regions. This high-resolution discrepancy at promoter NDRs between CUT&Tag and ChIP-seq profiles could potentially be explained by a local GC content and/or chromatin accessibility biases.

Altogether, our results highlight GC content and chromatin accessibility as important genomic features associated with divergent signals between ChIP-seq and CUT&Tag, which should be accounted for when selecting chromatin profiling approaches, analyzing and interpreting data, and drawing biological conclusions.

## Results

### Differential signal between ChIP-seq and CUT&Tag is associated with GC content and chromatin accessibility

To select functional regions of the genome associated with commonly studied histone marks for our comparative analysis, we use ChromHMM^24^ genomic annotations (**Fig. 1a**). We focused on three categories of genomic environments: active promoters, active gene bodies, and Polycomb-repressed domains (**Methods**). We strengthen the generalizability of our findings by analyzing public data from multiple labs and in two cell lines (K562 and MCF-7) for a range of histone modifications (H3K4me3, H3K27ac, H3K27me3 and H3K36me3; **Supp. Table 1**,**2**). We focused on histone marks associated with both narrow or broad genomic footprints and with both transcriptionally active or repressed chromatin. To identify regions relatively enriched in ChIP-seq vs CUT&Tag, we perform differential enrichment analysis with DESeq2^19^ comparing signals between the two chromatin profiling approaches on bins generated within the relevant genomic environment for each histone mark (**Fig 1a**; **Methods**). This strategy allows us to focus on genomic regions where we traditionally expect histone mark occupancy and isolate loci where ChIP-seq and CUT&Tag appear to disagree. In reconciling discrepant signals, we refrain from treating either assay as an absolute ground truth. Instead we assess independent metrics and complementary profiles (e.g., GC content, PRO-seq, DNase-seq) to identify genomic features associated with signal divergence and provide insight for the expected histone mark signal patterns in differentially-enriched regions (**Fig. 1a)**.

**Figure 1.**
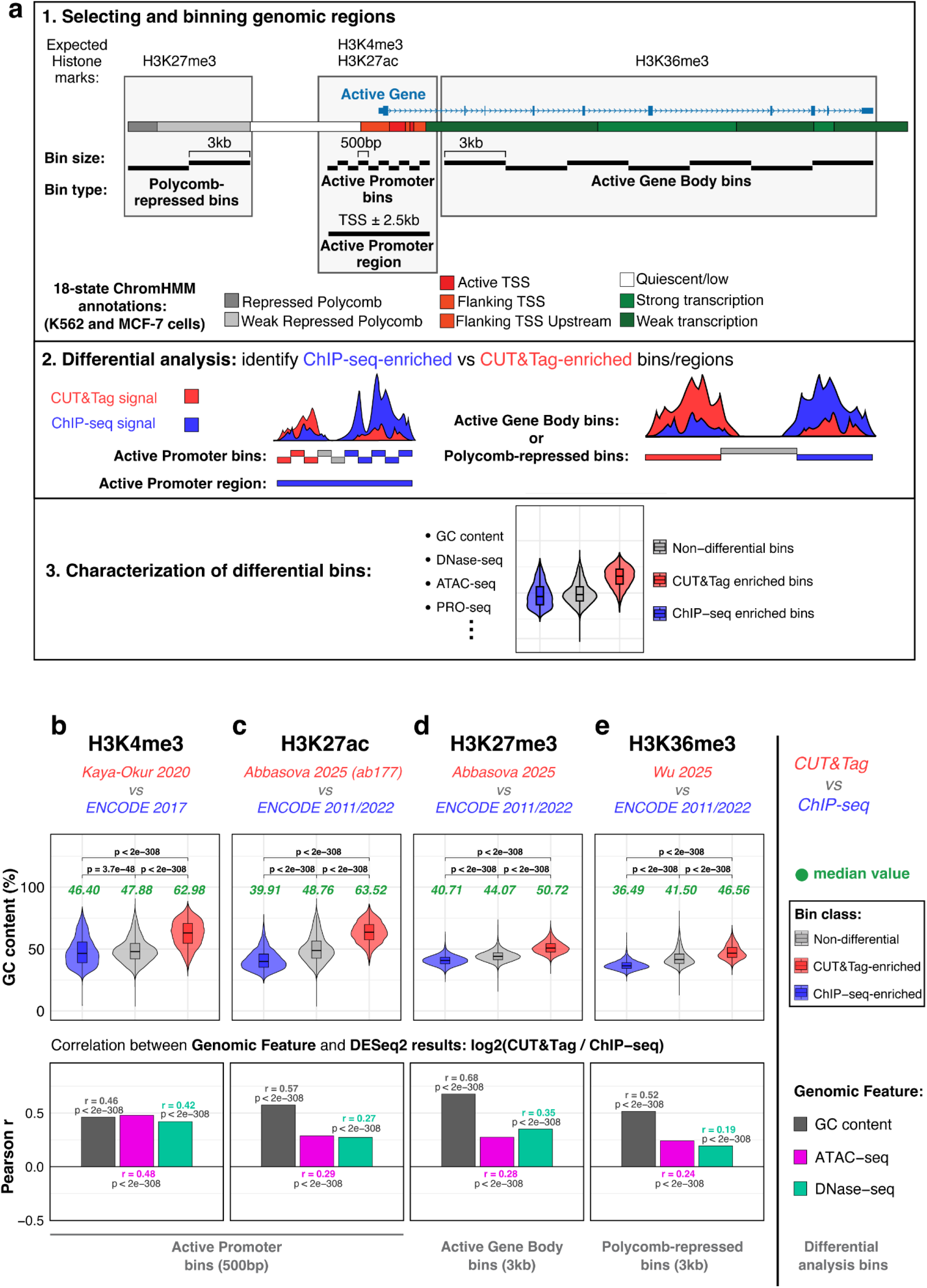
Differential signals between ChIP-seq and CUT&Tag are associated with GC content and chromatin accessibility. **a.** Schematic overview of differential analysis strategy for comparing ChIP-seq and CUT&Tag. **(1)** 18-state ChromHMM annotations for K562 and MCF-7 cell lines were used to select genomic regions associated with certain histone modification occupancy. Annotation track colors are the same as those used in the ChromHMM tracks^24^ **(2)** DEseq2^19^ was used to identify regions differentially-enriched between ChIP-seq and CUT&Tag. These designations denote regions of disagreement between the two assays, rather than loci uniquely captured by one method and not the other. For H3K4me3 and H3K27ac the differential analysis between ChIP-seq and CUT&Tag was performed on *Active Promoter regions* (active transcription start site (TSS) ± 2.5 kb; used in Fig. 3, **Supp. Fig. 1,6,9**). Differential analysis was also performed on *Active Promoter bins* generated by binning the 5kb Active Promoter region into 500bp bins (used in Fig. 1, **Supp. Fig. 1-6).** For H3K27me3 and H3K36me3, the differential analysis was conducted on Polycomb-repressed domains and active gene bodies, respectively, with the full regions segmented into 3 kb bins. See **Methods** and **Supp. Data 1**-**4** for more details. **(3)** GC content and orthogonal genomic signals such as DNase-seq and PRO-seq were used to characterize the differential regions between ChIP-seq and CUT&Tag. See **Methods** for additional details. **b-e. Top:** Violin plots showing the distribution of GC content (average GC% per bin) in ChIP-seq-enriched, CUT&Tag-enriched, and non-differential bin-sets. Differential bins are classified by DESeq2 (FDR < 0.05 and |log2FC| > 1.0). Median GC content is shown in green. Statistical differences in the distribution of GC content between the three bin subsets is calculated with a two-sided Wilcoxon rank−sum test, Benjamini−Hochberg corrected across the three comparisons within each panel. **Bottom:** the per-region or per-bin log2FC reported by DESeq2 was correlated against three genomic features: GC content (grey), ATAC-seq signal (magenta) and DNase-seq signal (teal). Bars show Pearson correlation coefficient r for each feature within each CUT&Tag vs ChIP-seq comparison; because fold change is defined as log2(CUT&Tag / ChIP-seq), positive r indicates that the feature is higher in regions enriched for CUT&Tag, negative r indicates it is higher in regions enriched for ChIP-seq. Correlations were computed across all tested bins/regions and the Benjamini–Hochberg adjusted p-value is corrected across all 72 correlations tested in **Supp. Fig. 2**). GEO/ENCODE accession numbers for datasets used in each comparison: **b** (CUT&Tag = GSM4308166 GSM4308167^8^; ChIP-seq = ENCSR668LDD), **c** (CUT&Tag = GSM8728355 GSM8728356^1^; ChIP-seq = ENCSR000AKP), **d** (CUT&Tag = GSM8728361 GSM8728362^1^; ChIP-seq = ENCSR000AKQ), **e** (CUT&Tag = GSM8343601 GSM8343602 GSM8343603 GSM8343604 GSM8343605^25^; ChIP-seq = ENCSR000AKR) (**Supp Table 1**).

A note on terminology and approach: throughout this analysis, *“ChIP-seq-enriched”* and *“CUT&Tag-enriched“* bins (or regions) refer to genomic loci identified by DESeq2 as relatively enriched in one assay compared to the other (**Methods**). These designations are based on the relative distribution of signal within each dataset for the selected regions, and therefore denote regions of disagreeing enrichment between the two assays, rather than loci uniquely captured by one method and not the other. To confirm that differentially-enriched regions represent loci with genuine signal over background rather than purely method-specific noise, we verified that regions enriched for either ChIP-seq or CUT&Tag show high concordance with the peaks called for their respective datasets. Specifically, for the set of selected ChromHMM annotated regions (namely, Active Gene Body bins, Polycomb-repressed bins, Active and Polycomb-repressed Promoter regions), we evaluated the overlap between CUT&Tag- and ChIP-seq-enriched bins/regions with each method’s peaks. As expected, CUT&Tag- and ChIP-seq-enriched loci largely show high overlap with peaks called for each corresponding chromatin profiling method (e.g., in K562, across all four histone marks, the overlap ranged from 75.7% - 98.6% for ChIP-seq peaks on ChIP-seq-enriched bins, and from 96.2% - 99.8% for CUT&Tag peaks on CUT&Tag-enriched bins; **Supp. Fig. 1**, **Supp. Data 5**; **Methods**). Note, potentially due to higher peak calling stringency or noisy environment definition in this cell line, ChIP-seq H3K27me3 peaks in MCF-7 cells showed a relatively lower degree of overlap with ChIP-seq-enriched Polycomb-repressed bins, at 38.1%. However this was still higher than the overlap with regions we do not expect to see H3K27me3 peaks (e.g., only 5.2% overlap with Active Gene Body bins), suggesting that these ChIP-seq-enriched Polycomb-repressed bins are not stemming from purely ChIP-seq noise (**Supp. Fig. 1**).

We next evaluated how two key genomic features, GC content and chromatin accessibility, associate with the differential signals between ChIP-seq and CUT&Tag within selected environments. For the active promoter associated histone marks H3K4me3 and H3K27ac, we first focused on the differential analysis performed on Active Promoter bins. Across all evaluated histone marks in K562 and MCF-7 cells, generated by at least 10 different groups, (a total of 14 CUT&Tag and 13 ChIP-seq datasets, with each dataset containing two or more biological replicates, leading to 24 unique comparisons; **Supp. Table 1**, **2**), GC content positively correlated with enrichment of CUT&Tag signal relative to ChIP-seq, with chromatin accessibility signal (DNase-seq and ATAC-seq) largely showing the same trend (**Fig 1b-e**, **Supp. Fig. 2-5**). For example, the median GC content of H3K4me3 CUT&Tag-enriched Active Promoter bins was 16.58% higher than the ChIP-seq-enriched bins (CUT&Tag = 62.98% while ChIP-seq = 46.40%; **Fig. 1b**; **Methods**). These differentially-enriched promoter bins also show distinct chromatin accessibility signal, with CUT&Tag-enriched bins showing higher median Z-scored DNase-seq signal compared to ChIP-seq-enriched bins (CUT&Tag-enriched = 0.69, ChIP-seq-enriched = -0.33; **Supp. Fig. 3b**; **Methods**). Similar accessibility related patterns were observed with ATAC-seq data (**Supp. Fig. 3c**). Genome browser tracks (and interactive session links) of example differential loci for all four histone marks across active promoters, active gene bodies and Polycomb-repressed domains in K562 cells, support the association of GC content and accessibility with the differential signals between the two chromatin profiling methods (**Supp. Fig. 6**; **Supp. Data 1**-**4**) provide coordinates for inspection of additional differential analysis results in the interactive browser sessions).

**Figure 2.**
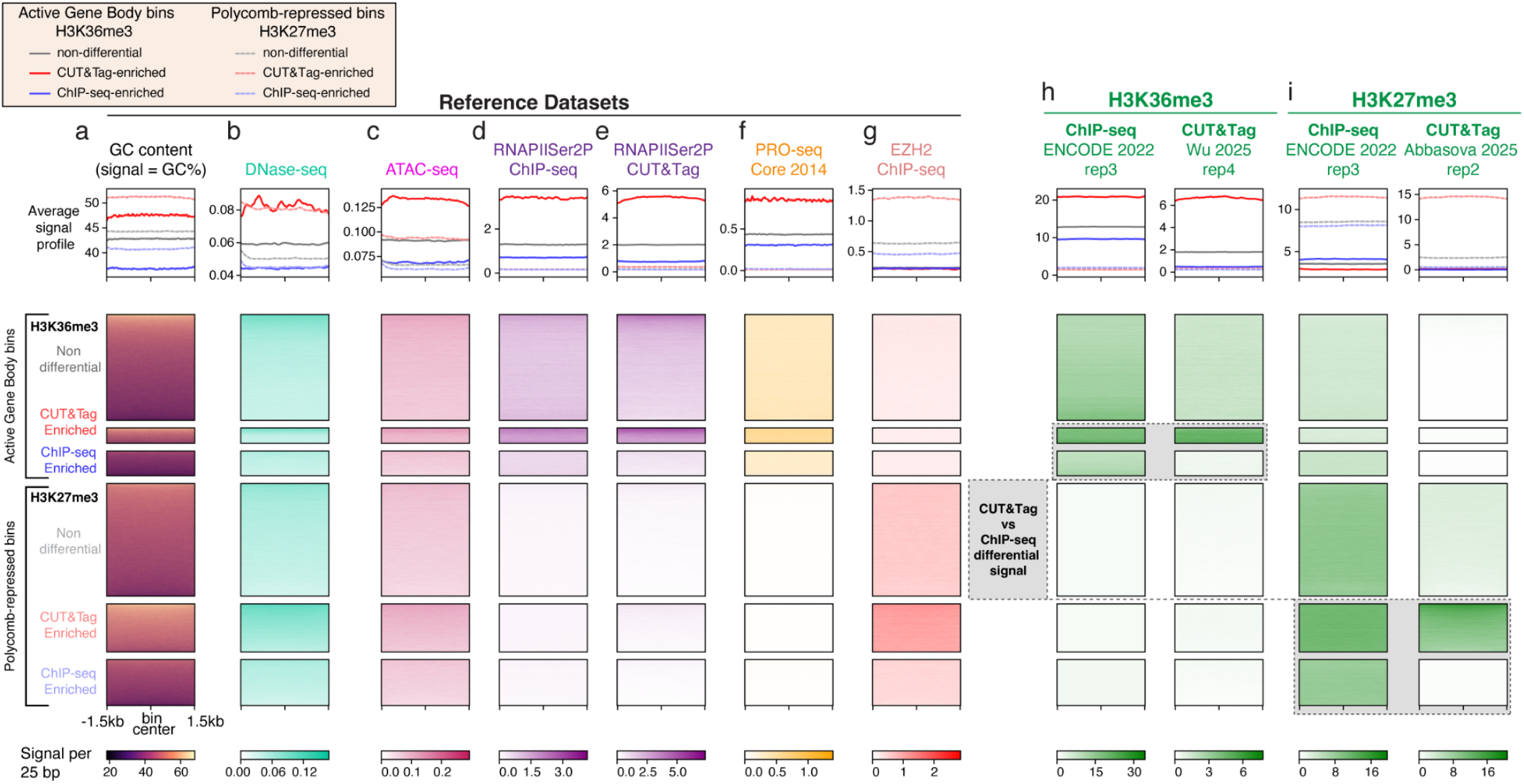
Regions showing limited H3K27me3 and H3K36me3 CUT&Tag signal compared to ChIP-seq exhibit low GC content and reduced chromatin accessibility. Heatmaps on differentially- and non-differentially-enriched Active Gene Body and Polycomb-repressed bin subsets identified by H3K36me3 and H3K27me3 differential analysis on each region set, respectively. All datasets are from K562 cells; heatmaps are computed by taking average signal in 25 bp bins across each 3 kb Active Gene Body or Polycomb-repressed bin, with the color ceiling set to each column’s 97th percentile to suppress outliers. Each bin subset is individually sorted by decreasing GC content, and is centred on the 3 kb bin (1.5 kb flanks). Above each heatmap column, a line plots represent the average signal (or in the case of GC content, the GC%) in 25 bp bins across the 3 kb bins for each subset. **a-g**. Reference datasets include GC content, ATAC-seq, PRO-seq, as well as RNAPIISer2P ChIP-seq and CUT&Tag and EZH2 ChIP-seq (see **Supp. Table 1,3** for accessions). **h,i**. ChIP-seq and CUT&Tag signal quantified from a representative sample. Grey box: ChIP-seq and CUT&Tag signal on differentially-enriched bins, classified by DESeq2 as having an FDR<0.05 and |Log2(CUT&Tag/ChIP-seq)| > 1.0 (**Methods**).

**Figure 3.**
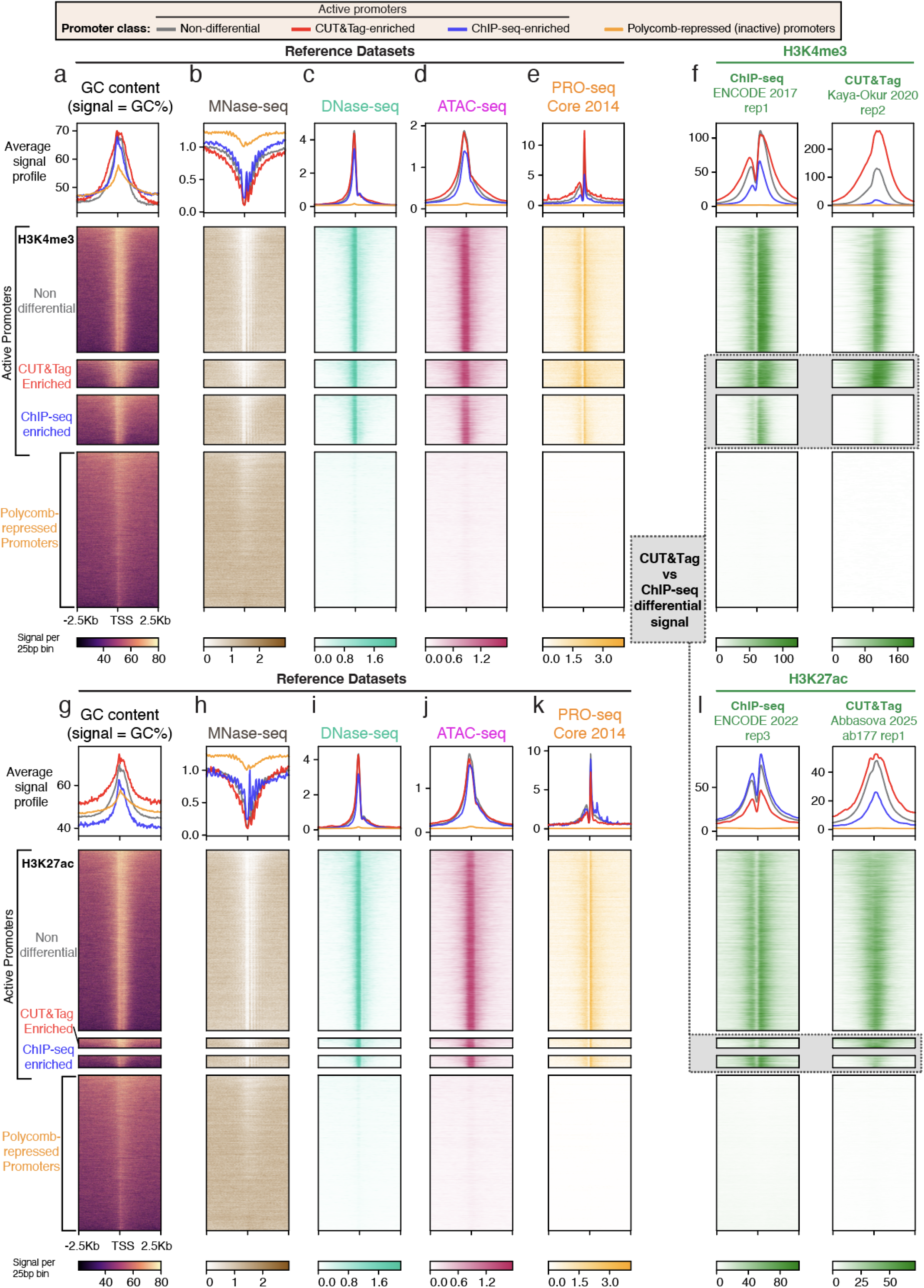
Active promoters showing differential H3K4me3 and H3K27ac ChIP-seq and CUT&Tag signals are associated with GC content and chromatin accessibility patterns. **a-l.** For H3K4me3 and H3K27ac a differential analysis between ChIP-seq and CUT&Tag was performed on Active Promoter regions (TSS ±2.5 kb). Heatmaps show signals quantified on ChIP-seq-enriched, CUT&Tag-enriched and non-differential Active Promoter regions as well as Polycomb-repressed Promoters Regions (used as a set of background regions representing silenced promoters). H3K4me3 and H3K27ac differentially-enriched promoter regions classified by DESeq2 as having an FDR<0.05 and |Log2(CUT&Tag/ChIP-seq)| > 1.0. (**Methods**) Data is from K562 cells; heatmaps are computed by taking average signal (or in the case of GC content, the GC%) in 25 bp bins across each Active or Polycomb-repressed Promoter Region (5kb), with color ceiling set to each column’s 97th percentile to suppress outliers. Each Active Promoter region subset is individually sorted by decreasing GC content and centered on the TSS with 2.5 kb flanks. Above each heatmap column, Average signal profile (metagene) plots represent the average signal in 25 bp bins across the 5 kb region for each promoter region subset. **a-e, g-k**. Reference datasets include GC content, MNase-seq, DNase-seq, ATAC-seq, PRO-seq (see **Supp. Table 1,3** for accessions). ChIP-seq and CUT&Tag signal for H3K4me3 (**f**) and H3K27ac (**l**) quantified from a representative sample. Grey box: ChIP-seq and CUT&Tag signal on differentially-enriched Active Promoter regions.

### Regions with limited H3K36me3 and H3K27me3 CUT&Tag signal relative to ChIP-seq are associated with low GC content and reduced chromatin accessibility

To further characterize the differentially-enriched regions identified above, we initially focused on the histone marks H3K36me3 and H3K27me3, which were analyzed on active gene bodies and Polycomb-repressed domains, respectively. Active Gene Body and Polycomb-repressed bins were generated by segmenting all selected regions into 3 kb bins, which were subsequently used to detect differentially-enriched loci within each environment (**Methods**). These histone marks mainly display broad signal patterns, are expected to be mostly mutually exclusive, and are associated with distinct genomic activity (H3K36me3: active transcription along gene body; H3K27me3: Polycomb-mediated transcriptional repression)^26^. To corroborate the expected functional characteristics of these selected genomic environments, we employed a range of orthogonal and complimentary reference datasets. The reference datasets, in K562 cells, associated with active transcription includes: PRO-seq (Core et al. 2014^27^ and Dastidar et al. 2023^28^), as well as ChIP-seq and CUT&Tag for RNA polymerase II subunit A (RNAPII) with phosphorylated serine 2 or 5 (RNAPIISer2P or RNAPIISer5P) (**Supp. Tables 1**-**3**). RNAPIISer2P maps the elongating form of RNAPII and RNAPIISer5P profiles RNAPII during initiation^29^. In line with our expectations regarding transcriptional elongation signal, PRO-seq and RNAPIISer2P ChIP-seq profiles showed stronger signal across all subsets of Active Gene Body bins compared to the Polycomb-repressed bin subsets (**Fig 2d**-**f**, **Supp. Fig. 7d**-**k, Supp. Fig. 8a**). The reference dataset associated with Polycomb-repressed regions includes ChIP-seq for the histone methyltransferase EZH2, a subunit of the Polycomb Repressive Complex 2 (PRC2), which is responsible for depositing H3K27me3^30–32^. Aside from RNAPII, we use ChIP-seq data for mapping of non-histone proteins, due to general CUT&Tag usage recommendations^7,8^ as well as data availability. As expected we observed higher EZH2 ChIP-seq signal across the Polycomb-repressed bins compared to the Active Gene Body bins (**Fig 2g**, **Supp. Fig. 7l**, **Supp. Fig. 8a**).

We next evaluated relative signal intensities of H3K36me3 and H3K27me3 ChIP-seq and CUT&Tag, in K562 cells, across the subsets of differentially-enriched bins (**Fig. 2, Supp. Fig. 7**). Between the different subsets of Active Gene Body bins, the CUT&Tag-enriched bins showed the highest median H3K36me3 signal in both ChIP-seq and CUT&Tag datasets (**Fig. 2h, Supp. Fig. 7m,n**). A parallel pattern was observed for the subsets of Polycomb-repressed bins, where the CUT&Tag-enriched subset showed the highest median H3K27me3 signal for both profiling methods (**Fig. 2i, Supp. Fig. 7o,p**). These CUT&Tag-enriched bins also demonstrated the highest relevant reference signals compared to the rest of the subsets. Specifically, for Active Gene Body bins, the CUT&Tag-enriched subset displayed the highest median signal across PRO-seq and the range of RNAPII localization profiles (**Fig. 2d-f, Supp. Fig. 7d-k**). For the set of Polycomb-repressed bins, the CUT&Tag-enriched subset showed the highest median EZH2 ChIP-seq signal (**Fig. 2g, Supp. Fig. 7l**). Notably, we observe elevated GC content and open chromatin signal at CUT&Tag-enriched bins relative to the other bin subsets (e.g., CUT&Tag-enriched Active Gene Body bins show a median GC content and DNase-seq signal of 46.6% and 0.07, respectively, whereas ChIP-seq-enriched bins show 36.5% and 0.05) (**Fig 2a-c, Supp. Fig. 7a-c**). The higher relevant reference dataset signal in CUT&Tag-enriched (higher GC) bins described above, aligns with previous studies linking GC-rich sequences with both RNAPII pausing^33^ and PRC2 recruitment^34^. Together, analysis of these multi-assay datasets robustly supports the presence of H3K36me3 and H3K27me3 signal at CUT&Tag-enriched bins within active gene bodies and Polycomb-repressed domains, respectively.

Notably, we observe a major disagreement between the two chromatin profiling methods in ChIP-seq-enriched bins, with CUT&Tag exhibiting an apparent reduction in sensitivity. Specifically, CUT&Tag shows a 6.07-fold reduction in median H3K36me3 signal, while ChIP-seq only shows a 1.35-fold reduction, when comparing CUT&Tag- to ChIP-seq-enriched Active Gene Body bins. This trend was mirrored when we compared the overlap of bin subsets with peaks called for each dataset, with CUT&Tag showing a reduction in overlap of 80.8% (from 96.2% to 15.4%) and ChIP-seq showing a reduction of only 6.6% (from 93.3% to 86.7%) (**Fig. 2h, Supp. Fig. 1f, 7m,n**). A similar trend was observed for H3K27me3 on Polycomb-repressed bins, with median CUT&Tag signal dropping 6.75-fold between the CUT&Tag- and ChIP-seq-enriched bins (peaks reducing overlap by 86.6%, from 99.8% to 13.2%), and ChIP-seq showing a drop of only 1.14-fold median signal (peaks reducing overlap by only 15.5%, from 91.2% to 75.7%) (**Fig 2i, Supp. Fig. 1e, 7o,p**).

To evaluate whether these apparent CUT&Tag-depleted loci are expected to have H3K36me3 or H3K27me3 signal, we employ the orthogonal and complementary K562 reference datasets discussed above. These reference datasets inform on active transcription and Polycomb localization, and thus can support the general presence of H3K36me3 or H3K27me3, respectively. As the deposition of H3K36me3 on gene body nucleosomes is a hallmark of actively elongating RNAPII^35,36^, we reference three relevant datasets, namely PRO-seq as well as ChIP-seq and CUT&Tag for the elongating form of RNAPII (RNAPIISer2P). Similarly, to evaluate Polycomb-repressed bins, we utilized ChIP-seq for EZH2, the primary methyltransferase responsible for H3K27me3 deposition^30–32^. We first compared the median signal of the relevant reference datasets (e.g., PRO-seq) between differentially-enriched bin subsets within the same genomic environment (e.g., CUT&Tag- and ChIP-seq-enriched Active Gene Body bins). Next, to gather evidence for whether a dataset shows signal in a particular bin subset that is greater than the dataset’s estimated background level, we compare signals across functionally distinct, largely mutually exclusive, genomic environments (e.g., we use Polycomb-repressed domains as background regions for signal expected on active gene bodies).

As mentioned above, in both genomic environments, we observe a reduction in the median signal of the relevant reference datasets in ChIP-seq-enriched bins compared to the CUT&Tag-enriched subset, which could potentially be explained by the difference in GC content between the differentially-enriched regions^33,34^ (**Fig 2, Supp Fig 7**-**9**). Yet, when compared to the background environment, the ChIP-seq-enriched bins still show a statistically significant higher median signal in relevant reference datasets. More specifically, the median PRO-seq and RNAPIISer2P ChIP-seq and CUT&Tag signal in ChIP-seq-enriched Active Gene Body bins remains significantly higher than in all Polycomb-repressed bin subsets (**Fig. 2d-f**, **Supp. Fig. 7f**,**h**,**j**,**k**, **Supp. Fig. 8a**). Likewise, the median EZH2 signal in ChIP-seq-enriched Polycomb-repressed bins is significantly higher than in any Active Gene Body subset (**Fig. 2g**, **Supp. Fig. 7l**). Similar results were observed using chromatin profiles from MCF-7 cells with PRO-seq and RNAPII ChIP-seq displaying a statistically significant higher median signal in CUT&Tag-depleted (ChIP-seq-enriched) Active Gene Body bins compared to the subsets in the Polycomb-repressed environment (**Supp Fig. 8b**, **9d**,**e**).

Altogether, the differential analysis comparing ChIP-seq and CUT&Tag for H3K36me3 and H3K27me3 on active gene bodies and Polycomb-repressed domains, respectively, identifies regions of low GC content and reduced chromatin accessibility signal with limited CUT&Tag signal and sensitivity compared to ChIP-seq. The presence of relevant reference signals at some of these regions above the estimated background suggests that these differential regions are not emanating only from ChIP-seq noise. Rather, this differential signal potentially represents a technical GC content and/or chromatin accessibility related bias in CUT&Tag, which may lead to a limited sensitivity in such regions.

### Differential H3K4me3 and H3K27ac ChIP-seq and CUT&Tag signals at active promoters are linked to GC content and chromatin accessibility patterns

Next, we conducted a differential analysis comparing ChIP-seq and CUT&Tag signals for H3K4me3 and H3K27ac using 500 bp Active Promoter bins (**Fig. 1b,c, Supp. Fig. 2-4**). We observed that GC content and chromatin accessibility are associated with high-resolution, *intra-promoter*, discrepancies between the two chromatin profiling methods. To evaluate the differential signals at the *inter-promoter* level, we performed a differential analysis using the full Active Promoter regions (TSS ± 2.5kb; described in **Fig. 1a**; **Methods**). This allowed us to identify Active Promoter regions that are relatively enriched in ChIP-seq vs CUT&Tag and vice versa. To independently evaluate the transcriptional activity of these selected promoters, we quantified signal from the reference K562 datasets PRO-seq, as well as ChIP-seq and CUT&Tag for various phosphorylation states of RNAPII, and EP300 ChIP-seq (EP300 is a histone acetyltransferase responsible for depositing H3K27ac and is associated with active promoters^37,38^). For a background control, we employed Polycomb-repressed (inactive) promoter regions based on ChromHMM annotations (**Methods)**. As expected, we observed strong enrichment of signal from these transcription related datasets across the selected Active Promoter regions, with little to no signal on the Polycomb-repressed Promoter regions (**Fig 3a, Supp. Fig. 10a,b**). In MCF-7 cells, we similarly used reference datasets (PRO-seq and ChIP-seq for RNAPII and EP300), showing patterns matching the ones observed in K562 (**Supp. Fig. 10c,d**).

For each subset of Active Promoter regions displaying differential enrichment between the two chromatin profiling methods, we examined the average signal profile of ChIP-seq and CUT&Tag H3K4me3 and H3K27ac (i.e., metagene plots for each promoter region subset; **Fig. 3**). For H3K4me3 datasets in both cell lines, the CUT&Tag-enriched Active Promoter regions showed strong signal in both ChIP-seq and CUT&Tag, corroborated by high overlap with called peaks (ChIP-seq peaks showed 97.5% and 98.1% overlap in K562 and MCF-7, respectively, and similarly CUT&Tag peaks showed 98.3% and 100.0%; **Fig. 3f**, **Supp Fig. 1c**; **Methods**). However, ChIP-seq-enriched promoters displayed a notable depletion of H3K4me3 CUT&Tag signal compared to ChIP-seq, which aligned with a reduction in overlapping CUT&Tag peaks (ChIP-seq peaks showed 95.4% and 82.4% overlap in K562 and MCF-7 cells, respectively, while CUT&Tag showed only 56.9% and 59.5%; **Fig. 3f**, **Supp Fig. 1c**; **Methods**). For Active Promoter regions with differentially-enriched H3K27ac signal, we observed a different pattern. While CUT&Tag shows the highest average signal profile and peak overlap with CUT&Tag-enriched promoter regions in both K562 and MCF-7 cells, ChIP-seq showed its highest signal and peak overlap with ChIP-seq-enriched and Non-differential promoter regions for K562 and MCF-7 cells, respectively (**Fig. 3l**, **Supp Fig. 1d**).

To investigate possible genomic features associated with differential ChIP-seq and CUT&Tag profiling of active promoter histone marks, we examined GC content and chromatin accessibility (DNase-seq and ATAC-seq) profiles across the differentially-enriched promoter region subsets. Average signal profiles for these genomic features across the distinct promoter region subsets reveals consistent differences in sequence and chromatin accessibility between the two enrichment classes, across all histone marks and cell lines. Specifically, CUT&Tag-enriched promoters on average show a higher, broader spike in GC content centered around the TSS, and higher chromatin accessibility signal (DNase-seq and ATAC-seq), compared to ChIP-seq-enriched promoters (**Fig 3a-d,g-j, Supp Fig. 10**). PRO-seq seems to show comparable signal on ChIP-seq- and CUT&Tag-enriched Active Promoter regions, across all histone marks and cell lines, except for H3K4me3 in K562 cells, where it shows less signal in ChIP-seg-enriched promoter regions compared to other subsets. However, the average signal profile intensity of these promoter regions is still notably higher than the background regions, suggesting that these promoters are still transcribed, yet are potentially less active. The general consistency of transcriptional signal across the differentially-enriched Active Promoter regions, alongside the patterns described regarding differing GC content and chromatin accessibility on these regions, suggests that sequence and accessibility based genomic features may be driving differential ChIP-seq and CUT&Tag signal on active promoters.

In addition to the discrepancies between ChIP-seq and CUT&Tag average signal intensity and peak overlap across the differentially-enriched Active Promoter regions, we also observed differences in the profile shape of H3K4me3 and H3K27ac signal between the two assays across all active promoters. Similar to observations made using an alternative transposase-based chromatin profiling method^23^, ChIP-seq and CUT&Tag show different spatial signal distribution around the nucleosome-depleted region (NDR; commonly located immediately upstream of active TSSs^22^). ChIP-seq and MNase-seq (a method for mapping nucleosome positioning^39^) both display a characteristic valley of signal intensity across the NDR, linked to the coordinated positioning of nucleosomes around the promoter TSS^22^, CUT&Tag shows a monotonic decrease in signal across this region, more reminiscent of the average signal pattern observed in GC content, DNase-seq, and ATAC-seq signal around the NDR region (**Fig. 3**, **Supp Fig. 10**). Together with the GC content and accessibility patterns described on differentially-enriched promoter regions, this discrepancy in signal shape between the two methods may also be linked to the hyper-accessibility and GC content structure of active promoters.

## Discussion

In this study, we employed a systematic comparative approach to detect and characterize ChIP-seq and CUT&Tag discrepancies. Our approach included using datasets generated by multiple groups and focusing on a range of well studied histone modifications profiled in two cell lines. We conduct differential analysis to identify loci relatively enriched to one method compared to the other, and leverage orthogonal and complementary datasets (e.g., PRO-seq, DNase-seq) to characterize regions of disagreement. By directly comparing signal between ChIP-seq and CUT&Tag (relative to each method’s own signal baseline) within previously annotated functional genomic regions, we can identify regions of discrepancy without fully basing our analysis on peak calling from individual samples, which may require sample specific optimizations such as the choice of peak caller and parameter, and threshold tuning. Note, we do use peak calling based analysis to corroborate our differential analysis based observations (**Supp. Fig. 1, 11**).

We identify GC content and chromatin accessibility as genomic features largely associated with differential signal between ChIP-seq and CUT&Tag across four key histone marks: H3K4me3, H3K27ac, H3K27me3 and H3K36me3. A recent study comparing ENCODE peaks to CUT&Tag peaks generated for H3K27ac and H3K27me3 reported that CUT&Tag-unique peaks show higher ATAC-seq signal compared to ENCODE-unique peaks^1^. We re-analyzed the overlap between these peaks and observed that ENCODE-unique peaks show a lower median GC content compared to CUT&Tag-unique peaks across all comparisons (i.e., across all peak callers, antibodies and biological replicates compared; **Supp. Fig. 11**). Specifically, H3K27ac CUT&Tag-unique peaks had a median GC content range of 50.8% – 57.6% across all comparisons, with ENCODE-unique peaks displaying a range of 45.2% – 47.5%. H3K27me3 CUT&Tag-unique peaks showed a median GC content range of 50.1% – 52.1% while ENCODE-unique had a range of 44.5% – 44.7% (**Supp. Fig. 11**). Thus, the observations made in this peaks-based analysis further support the association of differential ChIP-seq and CUT&Tag signals with GC content and chromatin accessibility.

Given that the CUT&Tag can be performed with different protocol parameters, such as the use of light fixation or choice of DNA isolation method, we selected CUT&Tag samples for H3K36me3 and H3K27me3 prepared using various conditions and compared the signal of each dataset on the differential bins previously identified within active gene bodies and Polycomb-repressed regions, respectively (**Supp. Fig. 12**). Across all conditions, the patterns we initially observed (see **Fig. 2**) were generally maintained. Specifically, CUT&Tag showed a stronger depletion of signal from CUT&Tag- to ChIP-seq-enriched regions, compared to ChIP-seq signal across these regions (**Supp. Fig. 12**). We did observe a potential slight recovery in CUT&Tag sensitivity in samples with light fixation. However, due to the limited number of replicates in each protocol condition and other challenges (i.e., low read depth in some samples), this was mostly inconclusive. Altogether, this analysis supports the generalizability of our identified differential regions as well as the notion that the discrepancies between CUT&Tag and ChIP-seq are associated with GC content and chromatin accessibility across various current CUT&Tag protocol conditions.

As previously observed, chromatin accessibility and genomic GC content can be associated with each other^40^, making it difficult to determine whether only one or both features are driving the associations of these genomic features with the divergent signals between ChIP-seq and CUT&Tag. Adding further complexity to the relationship between these features, the Tn5 transposase has been previously reported to have a preference for GC-rich regions in ATAC-seq data^41^. Based on our observations in this study, we propose that GC content and chromatin accessibility are each associated with differential signals between ChIP-seq and CUT&Tag, and that their effects are at least partly separable. Particularly for the broad marks H3K36me3 and H3K27me3, ChIP-seq vs CUT&Tag discrepancies frequently occur at regions lacking characteristic DNAse-seq and ATAC-seq signal spikes and instead track more closely with the underlying GC content (**Supp. Fig. 6c,d**). Additionally, some of the comparisons between K562 and MCF-7 CUT&Tag and ChIP-seq data for H3K27ac show a negative correlation between chromatin accessibility and CUT&Tag-enriched signal, while GC content showed a positive correlation for all comparisons made in this study (**Supp. Fig. 2, 4b,c**). Finally, MA- (minus vs average) plots and volcano plots depicting the differential analysis results were colored by the GC content, DNase-seq or ATAC-seq signal, to allow visual inspection of the trending between genomic features and differential signal (**Supp. Fig. 3-5**). From these visualizations we note that chromatin accessibility signals did not show fully overlapping trends with GC content. Altogether, these observations indicate that GC content and chromatin accessibility can associate with signal disparity between the two methods at least partly independent of each other, suggesting that their associations may each be arising from distinct technical biases or sensitivities.

Several recent studies report discrepancies between ChIP-seq and CUT&Tag, with some proposing novel binding patterns for histone marks and transcription factors that disagree with, or alter, previously established ChIP-seq-based results^15–18^. Our findings identify potential sources of technical bias/sensitivity that may contribute to differential signals between these two methods, and some of the discrepant conclusions mentioned above. Overall, such contradicting results alongside our main observations underscore that apparent disagreements between chromatin-profiling assays may arise from method-specific biases rather than biology, and that accounting for such biases is essential when designing experiments, analyzing data, and drawing biological conclusions from any single approach.

Our study has several limitations. The use of previously defined ChromHMM regions (with annotations based on ChIP-seq data) for our differential analysis inherently restricts the ability to detect CUT&Tag-unique differential signals that may occur outside of these selected loci. However, our re-analysis of genome-wide peaks generated by a recent study^1^ also links these disagreeing signal detection to GC content, in addition to chromatin accessibility, suggesting that our main conclusions are not dependent on this limitation (**Supp. Fig. 11**). We also note that while DESeq2^19^ provides a useful framework to compare count data such as ChIP-seq and CUT&Tag signal in genomic bins, it was not specifically designed to compare orthogonal chromatin-profiling approaches. Thus, differences in library composition, signal-to-noise ratios, dynamic range, and normalization between the two assays may result in differential calls that are approximate, and thus our approach may not perfectly partition the signal discrepancies between the two methods.

Altogether, the observations presented in this analysis highlight that studies focused on profiling the intensity and structure of histone modification occupancy at genomic regions such as promoters, gene bodies and Polycomb-repressed domains, may be sensitive to the biases of CUT&Tag or ChIP-seq to GC content and chromatin accessibility. Further, our study demonstrates the power of integrating orthogonal measurements of relevant omic readouts to corroborate or question the results obtained by each chromatin profiling method.

## Methods

### Data acquisition and processing

Published sequencing data were downloaded from the Sequence Read Archive with prefetch and fasterq-dump --split-files (sra-tools 3.2.1, https://github.com/ncbi/sra-tools) or, for ENCODE^42–44^ (https://www.encodeproject.org/) libraries, downloaded directly by file accession from ENCODE. All libraries (ChIP-seq, CUT&Tag and ATAC-seq) were processed through a pipeline (**GitHub:** https://github.com/edumodolo/modolo_et_al_2026_CUTnTag_ChIPseq_analysis). Technical replicates were concatenated before trimming; biological replicates were kept separate for all ChIP-seq and CUT&Tag samples, so that replicate-level variability is preserved in the differential analysis.

Adapters and low-quality bases were removed with fastp^45^ at default settings, with --detect_adapter_for_pe enabled for paired-end libraries. Reads were aligned with bowtie2^46^ to hg38. CUT&Tag paired-end libraries were aligned with `--local --very-sensitive --no-mixed --no-discordant --phred33 -I10 -X 700` (as recommended in^8^; https://dx.doi.org/10.17504/protocols.io.bjk2kkye), while ChIP-seq single-end libraries used `--local --very-sensitive --phred33`, and ATAC-seq samples used `-X 2000`. Alignments were filtered with samtools^47^ (removing unmapped reads, reads with an unmapped mate, secondary alignments, vendor-QC failures and flagged duplicates) at a MAPQ threshold of 20 for ChIP-seq and CUT&Tag and 30 for ATAC-seq. Paired-end libraries additionally filtered for properly paired reads. Duplicates were removed with `samtools markdup -r`, and BAM files were restricted to chr1–22, chrX and chrY (discarding chrM, unplaced contigs, random scaffolds and alternative haplotypes). ENCODE hg38 blacklist (v2)^48^ was subtracted from the final BAM file with bedtools^49^ intersect -v.

Coverage tracks were generated with deepTools^50^ bamCoverage at 1 bp resolution. Coverage is fragment-based throughout (i.e., paired-end reads were extended to their mate specified fragment size and single-end reads were extended to a fixed 200 bp). Per-library (biological replicate) tracks were left unnormalized (raw coverage). ATAC-seq biological replicates were merged into a single CPM-normalized track per cell line to generate the reference datasets used throughout this analysis.

HOMER v5.1^51^ (`makeTagDirectory -checkGC -fragLength 200`), was used to quantify nucleotide frequency around 5′ read ends and fragment GC distribution against the hg38 background in 200 bp windows (available on GitHub: https://github.com/edumodolo/modolo_et_al_2026_CUTnTag_ChIPseq_analysis). The pipeline compiles a per-library metrics table (raw read count, overall alignment rate, fraction of reads passing filtering, duplication rate, mitochondrial fraction and final mapped fragment count) reported in **Supp. Table 2**.

### Reference datasets

hg38 GC content was downloaded from UCSC Genome Browser^52^ as a 5 bp resolution BigWig, (hg19 wig file was converted to BigWig with wigToBigWig^53^ for **Supp. Fig. 11**). DNase-seq, MNase-seq, and EZH2, EP300 and RNA polymerase II (RNAPII, RNAPIISer5P, RNAPIISer2P) ChIP-seq signal tracks were obtained from ENCODE^42–44^ (https://www.encodeproject.org/). The hg19 MNase-seq and PRO-seq^27,28^ tracks were lifted to hg38. Stranded PRO-seq bigWigs were combined into a single unstranded track with deepTools bigwigCompare at 25 bp bins. (Accessions for all datasets are listed in **Supp. Tables 1,3**).

### Differential analysis

#### Selecting Region sets (**Fig. 1a**)

Analysis regions were built per cell line from the ENCODE 18-state ChromHMM models^54^ for K562 (ENCFF963KIA) and MCF-7 (ENCFF985EWD). A gene (canonical gene-body annotations, hg38) was defined as active when at least 20% of its body was covered by the Tx or TxWk states (`bedtools coverage`) and its TSS ± 1 kb overlapped an active-promoter state (TssA, TssFlnk, TssFlnkU or TssFlnkD). Four region sets were generated: (i) **Active Promoter regions**, TSS ± 2.5 kb (5 kb) of active genes, used both as whole windows and tiled into 500 bp **Active Promoter bins**; (ii) **Active Gene Body bins**, merged active gene bodies tiled into 3 kb bins with TSS ± 3 kb excluded; (iii) **Polycomb-repressed bins**, ReprPC/ReprPCWk domains tiled into 3 kb bins, with the bivalent states (TssBiv, EnhBiv) retained only where they fall inside a core ReprPC/ReprPCWk domain; and (iv) **Polycomb-repressed Promoter regions**, a negative-control set of promoters (TSS ± 2.5 kb) of genes whose TSS ± 1 kb overlaps a ReprPC domain, does not overlap any active-promoter state, and whose body carries no Tx/TxWk signal. Tiling used `bedtools makewindows`, retaining only tiles of the full bin width; all sets were restricted to chr1–22, X, Y, blacklist-subtracted and de-duplicated.

#### Signal quantification for features

Per-bin GC content and chromatin accessibility were quantified as the mean bigWig value over each interval with deepTools multiBigwigSummary BED-file --outRawCounts, computed once per cell line × region set so that every comparison over the same bins reads identical feature values.

#### Fragment counting

Count matrices for differential enrichment analysis were generated by counting fragments per bin directly from the filtered, deduplicated BAMs with deepTools multiBamSummary BED-file

`--outRawCounts` (deepTools 3.5.6), batched by cell line × region set × library layout. Paired-end batches were counted with `--extendReads --samFlagInclude 64` (counting first-in-pair reads only). Single-end batches were counted with `--extendReads 200`.

#### Differential enrichment

Count matrices were assembled by sample name and analysed with DESeq2^55^ (R 4.3, Bioconductor 3.18) using the design ∼ condition with ChIP-seq as the reference level and `fitType = “local“`. Bins with zero counts across all samples were removed prior to testing. The tested contrast was CUT&Tag versus ChIP-seq, so a positive log2 fold change denotes CUT&Tag enrichment and a negative value ChIP-seq enrichment. Bins or regions were classified as CUT&Tag-enriched (log2FC > 1.0 and FDR < 0.05), ChIP-seq-enriched (log2FC < −1.0 and FDR < 0.05) or non-differential; the same pair of thresholds (|log2FC| > 1.0, Benjamini–Hochberg FDR < 0.05) is used throughout the manuscript. Each CUT&Tag sample group was compared against every independent ChIP-seq source available for that histone mark and cell line, giving 24 comparisons in total (**Supp. Figs. 3-5**, **Supp. Tables 1, 2**). For genome browser inspection, bins/regions were additionally ranked by a signed π-score (log2FC × −log10(FDR), with FDR floored at 1 × 10^−300^), from which representative CUT&Tag-enriched, ChIP-seq-enriched and median non-differential active promoter regions were selected (**Supp. Fig. 6a**,**b, Supp. Data 1-4**). Differential-enrichment status-color bigBed custom tracks were produced with bedToBigBed^53^ and displayed on genome browser (**Supp. Fig. 6**).

#### Differential-analysis figures (Fig. 1d-e, Supp. Figs. 3-5)

Violin, volcano and MA panels were drawn in R with ggplot2^56^. Violin plots show the distribution of a genomic feature across the three differential classes, with an inset boxplot and the group median annotated. GC content is plotted and colored as mean GC% within a region/bin, presented on a fixed scale of 30–75% with no standardization. Chromatin accessibility (DNase-seq, ATAC-seq) is summarized as log2(mean signal + 1) and converted to a Z-score using the mean and standard deviation computed within each cell line × region set (e.g., Active Promoter bins standardized against Active Promoter bins only), so that intervals of different size and chromatin class are never pooled; the color range is ±3 SD with values beyond the range clamped. In the volcano plots −log10(FDR) is capped at 300; MA plots use log2(baseMean + 1). Statistical comparisons between the three differential classes within a violin plot panel used two-sided Wilcoxon rank-sum tests on all three pairwise contrasts, Benjamini–Hochberg-corrected within each panel.

#### Feature–log2FC correlation (Supp. **Fig. 2**)

The association between differential signal and each genomic feature was quantified as the Pearson correlation between the per-bin log2FC reported by DESeq2 and the per-bin feature value (i.e., mean GC%, log2(ATAC-seq + 1) and log2(DNase-seq + 1)), using cor.test (H₀: ρ = 0) over all tested bins. Because log2FC is log2(CUT&Tag / ChIP-seq), a positive r indicates that the feature is higher in CUT&Tag-enriched regions and a negative r that it is higher in ChIP-seq-enriched regions. Correlations were computed on the binned region sets (500 bp Active Promoter bins for the promoter marks, 3 kb bins for the broad marks). All correlations for the complete comparison set were computed in a single pass and the Benjamini–Hochberg correction was applied once across all 72 tests.

### Peak calling and peak overlap analysis (Supp. Fig. 1)

#### Peak calling

Peaks were called with MACS2^57^ 2.2.9.1. Common parameters between ChIP-seq and CUT&Tag were `-g hs -q 1e-5 --nolambda --keep-dup all --nomodel`. Narrow mode was used for the promoter marks (H3K4me3, H3K27ac) and `--broad --broad-cutoff 0.1` for the broad marks (H3K27me3, H3K36me3). The only deliberate difference between methods was the input format: ChIP-seq libraries, which were single-end, were read as `-f BAM` with a fixed `--extsize 200`, whereas CUT&Tag libraries, which are always paired-end, were read as `-f BAMPE`. Peaks called in each biological replicate were merged together into a master peak set for overlap analysis.

#### Overlap quantification

Each differential analysis results table was split into ChIP-seq-enriched, CUT&Tag-enriched and non-differential bins/regions (using thresholds described above), and the fraction of bins/regions in each class overlapping a merged peak set was counted with `bedtools intersect -u` at a minimum overlap of 1 bp. Each panel additionally carries a negative-control region set, where peaks for a given dataset are not usually expected.

#### Re-analysis of published CUT&Tag versus ENCODE peak sets (Supp. **Fig. 11**)

We re-analysed the H3K27ac and H3K27me3 CUT&Tag peak calls of Abbasova et al.^1^ against matched ENCODE ChIP-seq peaks, working in hg19 throughout. Published CUT&Tag peak files (MACS2 narrowPeak/broadPeak and SEACR stringent calls, per antibody and biological replicate) were retrieved from the authors’ CyVerse repository, and ENCODE K562 replicated peaks were downloaded from ENCODE for H3K27ac (ENCFF044JNJ) and H3K27me3 (ENCFF001SZF). All peak files were reduced to sorted, de-duplicated three-column BED intervals. Three subsets were defined per CUT&Tag sample with bedtools intersect: CUT&Tag-unique (CUT&Tag peaks with no overlapping ENCODE peak, -v), ChIP-seq-unique (ENCODE peaks with no overlapping CUT&Tag peak, -v) and shared (ENCODE peaks with at least 1 bp overlap, -u; the shared set is therefore defined on ENCODE peak boundaries). GC content per peak (mean GC content bigWig signal within every peak) was extracted from an hg19 gc5Base bigWig with `multiBigwigSummary BED-file --outRawCounts`. Subsets were compared by two-sided Wilcoxon rank-sum tests over the three pairs of subsets, Benjamini–Hochberg-corrected within each peak caller; the CUT&Tag-unique versus ChIP-seq-unique comparison is annotated on the figure.

### Signal quantification: heatmaps and violin plots

#### Region groups

For each heatmap or violin plot, data was quantified on the relevant set of region groups identified as ChIP-seq-enriched, CUT&Tag-enriched and non-differential by DESeq2 (FDR < 0.05 and |log2FC| > 1.0), as well as the matching negative-control set. Polycomb-repressed Promoter regions were used as the negative control region set for quantification of active promoter mark signal and reference datasets. Polycomb-repressed bins were used as the control for H3K36me3 and Active Gene Body bins for H3K27me3, exploiting the mostly mutual exclusivity of these two marks. The active promoter mark figures use the differential analysis results from the full Active Promoter region (TSS ± 2.5 kb) and the broad-mark figures use the 3 kb bin results.

#### Heatmaps and average signal profiles (**Figs. 2,3**, Supp. **Figs. 8, 10, 12**)

Signal matrices were built with deepTools `computeMatrix reference-point --binSize 25 --missingDataAsZero`, anchored at the TSS with ±2.5 kb flanks for the promoter marks and at the bin center with ±1.5 kb flanks for the 3 kb broad-mark bins, and rendered with `plotHeatmap --plotType lines`, which draws the average signal profile (metagene) for every region group above each heatmap column. Region groups were sorted independently, in descending order using the mean value per region of the first column (GC content). Every column carries its own independently scaled color scale and profile y-axis, so that tracks with a greater dynamic range cannot flatten the others. The color ceiling is set to the 97th percentile of each column (top 3% clipped), the color floor is 0 except for the GC column (which uses its 3rd percentile). Heatmaps were drawn with a fixed colour identity per data type (GC content, magma; DNase-seq, white→teal; ATAC-seq, white→magenta; all RNA polymerase II forms, white→purple; PRO-seq, white→orange; EP300, white→blue; EZH2, white→red; histone marks, white→green). Main-text figures are column subsets of the corresponding supplementary matrix, extracted with `computeMatrixOperations subset`, so that both figures are drawn from identical underlying values.

#### Violin plots (Supp. **Figs. 7**, 9)

For each bin/region group, the mean signal for every track over every interval was quantified with `multiBigwigSummary BED-file --outRawCounts` and plotted as log2(mean signal + 1); GC content is plotted untransformed as mean GC% per interval. Statistical testing used two-sided Wilcoxon rank-sum tests over all pairs of region groups present in a panel (i.e., fifteen pairs across the 6 different groups), with Benjamini–Hochberg correction applied within each panel.

#### Depth-matched comparison of CUT&Tag protocol variants (Supp. **Fig. 12**)

To compare CUT&Tag protocol conditions against one another, every CUT&Tag library was first subsampled to a common read depth of 1,000,000 fragments with `samtools view --subsample` at a fixed seed (`--subsample-seed 42`), using a fraction of 10⁶ divided by the library’s own fragment count. Fragment counts were taken as the number of first-in-pair alignments (`samtools view -c -f 64 -F 0×900`). ChIP-seq was not downsampled and is shown at original read depth for reference only. Heatmap quantification for this comparison uses the same per-column scaling as heatmaps described above, and H3K36me3/H3K27me3 differential regions are defined by the comparison completed in **Fig 1d,e**, and shown in **Fig. 2**.

## Supporting information

Supplemental Figure 3 (high resolution)

Supplemental Figure 4 (high resolution)

Supplemental Figure 5 (high resolution)

Supplemental Figure 7 (high resolution)

Supplemental Figure 10 (high resolution)

## Software and code availability

All processing and analysis scripts, together with the environment specifications, are available on GitHub: https://github.com/edumodolo/modolo_et_al_2026_CUTnTag_ChIPseq_analysis.

## Acknowledgements

We thank R. Wachs for her help with the illustrations. E. Modolo was supported in part by an institutional award to the UCSD Genetics Training Program from the National Institute for General Medical Sciences (grant no. T32 GM145427).

