## Supplemental Figures for "Discrepancies between ChIP-seq and CUT&Tag histone mark profiles are explained by GC content and chromatin accessibility"

### Supplementary Tables and Data:

**Supp. Table 1:**

Meta data collected for all ChIP-seq, CUT&Tag, and ATAC-seq data that was downloaded and processed for this analysis.

**Supp. Table 2:**

Post-processing metrics for ChIP-seq, CUT&Tag, and ATAC-seq samples processed for manuscript (technical replicates merged)

**Supp. Table 3:**

Meta data and download link for reference datasets downloaded and used for this manuscript

**Supp. Data 1-4**

csv files for example DEseq2 results which provide π-score ranked coordinates for inspection of additional differential regions using the interactive browser sessions.

**Supp. Data 5**

Data from Macs2 called peaks overlap with region subsets shown in (**Supp. Fig. 1**)

### Supplementary Figures:

*(Supplementary figures 3,4,5,7,10 attached as high resolution PDFs)*

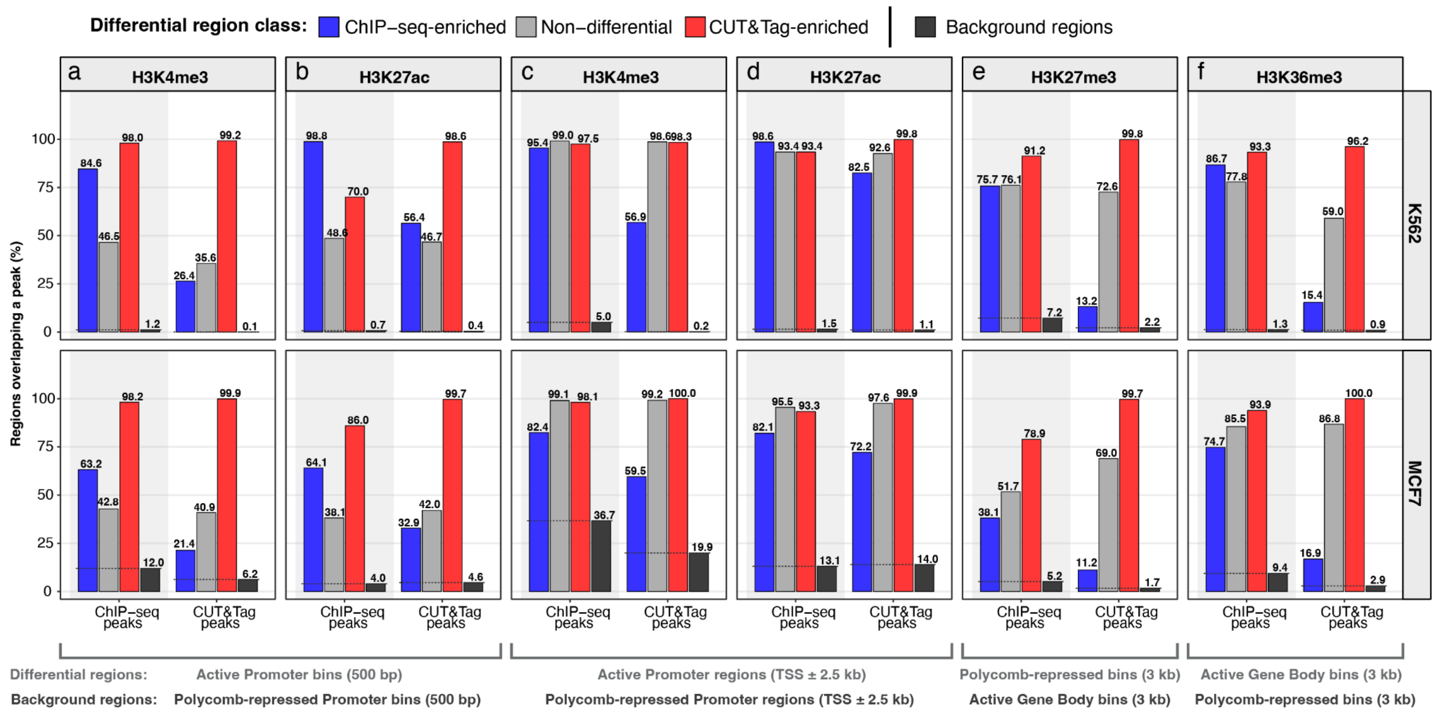


***Supplementary Figure 1. ChIP-seq and CUT&Tag MACS2 peaks overlap with differential and non-differential bin subsets.*** Peaks called for ChIP-seq and CUT&Tag with representative datasets for H3K4me3, H3K27ac, H3K27me3 and H3K36me3 using MACS2 (**Methods**) were overlapped with differential (ChIP-seq-enriched - blue bars, CUT&Tag-enriched - red bars), non-differential - light grey bars, and relevant background regions - dark grey bars. Percentage of differential or non-differential regions/bins overlapping a peak was plotted.

Differential regions were identified by comparing ChIP-seq and CUT&Tag signal with DESeq2 (FDR < 0.05, |log2FC| > 1.0) (see **Fig 1a**; **Methods).**

**a-d**) H3K4me3 and H3K27ac peak overlaps use differential analysis results from comparisons on Active Promoter Bins (500 bp) **a,b** and full Active Promoter regions (TSS ±2.5 kb) **c,d**, with Polycomb-repressed (inactive) Promoter bins and regions used as relevant background regions, respectively.

**b**) H3K27me3 and H3K36me3 peak overlaps use differential analysis results from comparisons on Polycomb-repressed bins and Active Gene Body bins, respectively.

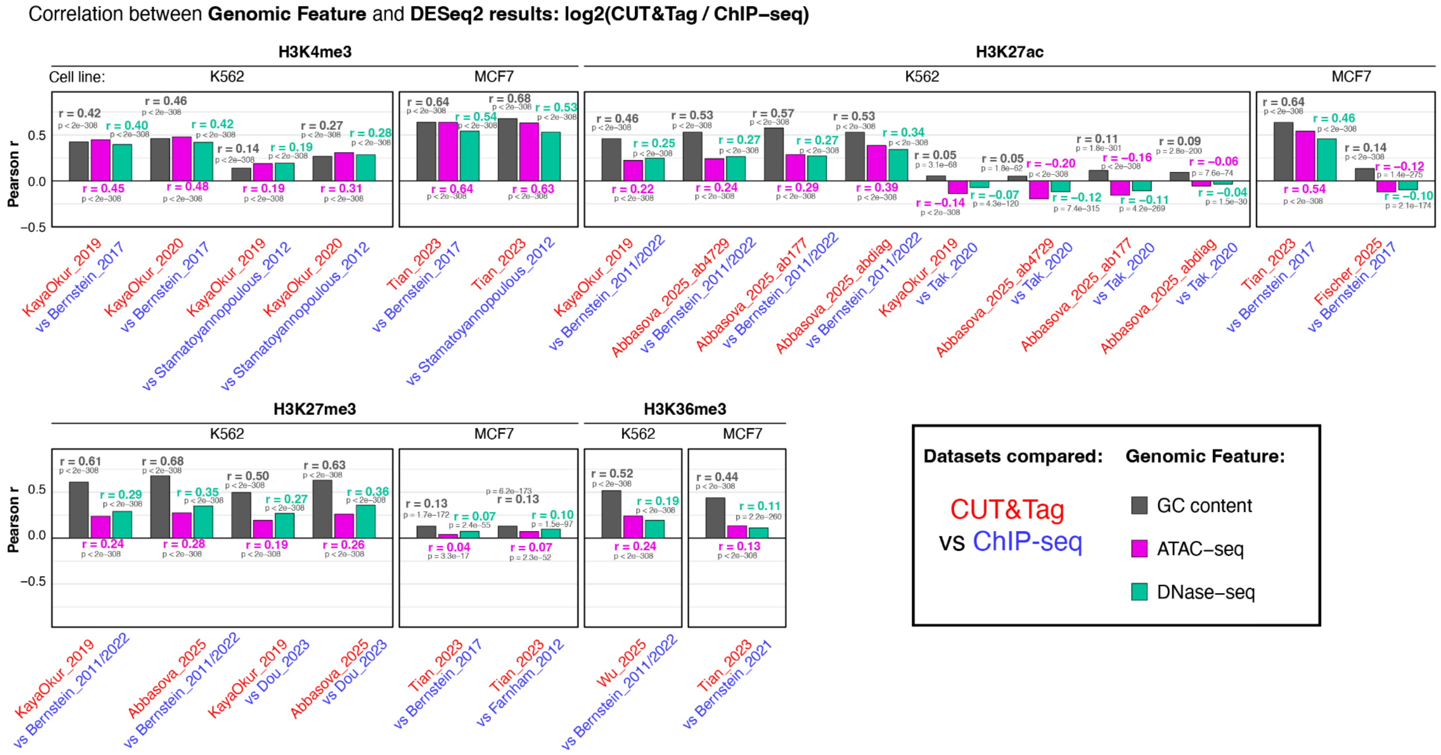


**Supplementary Figure 2.** ***Correlation of GC content and chromatin accessibility signal with ChIP-seq vs CUT&Tag differential analysis results***

The per-bin log2FC reported by DESeq2 was correlated against the per-bin signal of three genomic features: GC content (grey), ATAC-seq signal (magenta) and DNase-seq signal (teal). Bars show Pearson correlation coefficient r for each feature within each CUT&Tag vs ChIP-seq comparison; because log2FC is defined as log2(CUT&Tag / ChIP-seq), positive r indicates that the feature is higher in bins enriched for CUT&Tag, negative r indicates it is higher in bins enriched for ChIP-seq. Correlations were computed across all tested bins/regions and the Benjamini–Hochberg adjusted p-value is corrected across all 72 correlations tested in this figure.

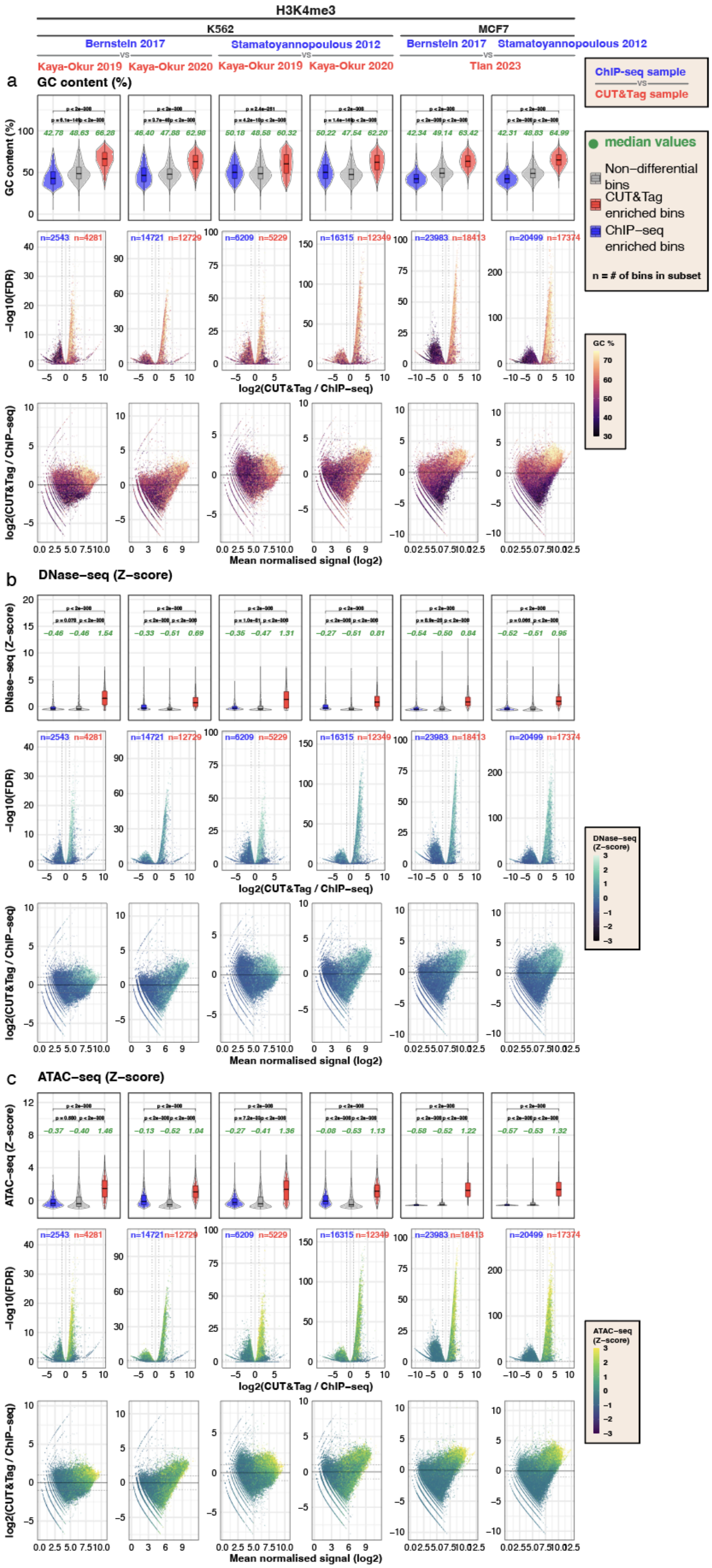

H3K4me3
K562
MCF7
Bernstein 2017
Stamatoyannopoulous 2012
vs
Kaya-Okur 2019
Kaya-Okur 2020
Tian 2023
ChIP-seq sample
CUT&Tag sample
a
GC content (%)
median values
Non−differential bins
CUT&Tag enriched bins
ChIP−seq enriched bins
n = # of bins in subset
GC %
−log10(FDR)
log2(CUT&Tag / ChIP−seq)
Mean normalised signal (log2)
b
DNase−seq (Z−score)
DNase−seq (Z−score)
c
ATAC−seq (Z−score)
ATAC−seq (Z−score)

*(high resolution images attached separately as PDF)*

**Supplementary Figure 3.** ***Differential signals between CUT&Tag and ChIP-seq for H3K4me3 are associated with GC content and chromatin accessibility across multiple datasets in K562 and MCF-7 cell lines*.**

**a-c. Top row:** Violin plots showing the distribution of GC content (**a**) Z-scored DNase-seq signal (**b**) or Z-scored ATAC-seq signal (**c**) in ChIP-seq-enriched, CUT&Tag-enriched, and non-differential Active Promoter bin subsets. Additional metadata and general processing metrics for CUT&Tag (red) and ChIP-seq (blue) samples compared are available in **Supp Table 1,2**. Median value is shown in green. Differential bins are classified by DESeq2 (FDR < 0.05 and |log2FC| > 1.0). Adjusted P-value for the differences in distribution of GC content or Z-scored DNase-seq or Z-scored ATAC-seq signal between the three bin subsets is calculated with a two-sided Wilcoxon rank−sum test, Benjamini−Hochberg corrected across the three comparisons within each panel. **Middle and Bottom rows**: Volcano plots and MA-plots for each CUT&Tag vs ChIP-seq comparison with each bin colored by its GC content, Z-scored DNase-seq or Z-scored ATAC-seq signal. Z-score signal for DNase-seq and ATAC-seq was calculated for each cell type independently.

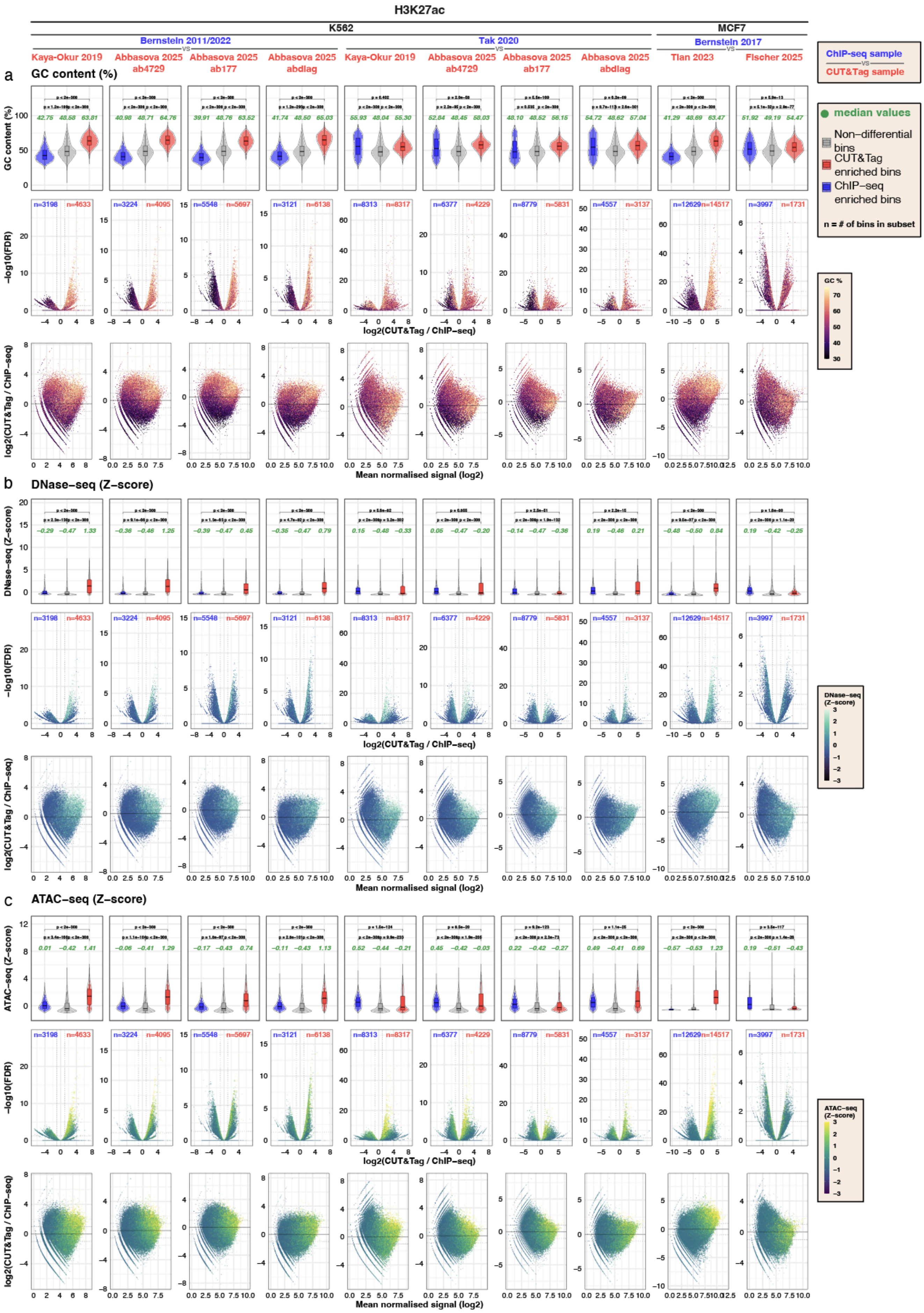
H3K27ac
K562
MCF7
Bernstein 2011/2022
Tak 2020
Bernstein 2017
vs
Kaya-Okur 2019
Abbasova 2025 ab4729
Abbasova 2025 ab177
Abbasova 2025 abdiag
Tian 2023
Fischer 2025
ChIP-seq sample
CUT&Tag sample
a
GC content (%)
b
DNase-seq (Z-score)
c
ATAC-seq (Z-score)
−log10(FDR)
log2(CUT&Tag / ChIP-seq)
Mean normalised signal (log2)
median values
Non-differential bins
CUT&Tag enriched bins
ChIP-seq enriched bins
n = # of bins in subset
GC %
DNase-seq (Z-score)
ATAC-seq (Z-score)

*(high resolution images attached separately as PDF)*

**Supplementary Figure 4.** ***Differential signals between CUT&Tag and ChIP-seq for H3K27ac are associated with GC content and partly associated with chromatin accessibility across multiple datasets in K562 and MCF-7 cell lines*.**

**a-c. Top row:** Violin plots showing the distribution of GC content (**a**) Z-scored DNase-seq signal (**b**) or Z-scored ATAC-seq signal (**c**) in ChIP-seq-enriched, CUT&Tag-enriched, and non-differential Active Promoter bin subsets. Additional metadata and general processing metrics for CUT&Tag (red) and ChIP-seq (blue) samples compared are available in **Supp Table 1,2**. Median value is shown in green. Differential bins are classified by DESeq2 as having (FDR $< 0.05$ and $|\log2FC| > 1.0$). Adjusted P-value for the differences in distribution of GC content or Z-scored DNase-seq or Z-scored ATAC-seq signal between the three bin subsets is calculated with a two-sided Wilcoxon rank−sum test, Benjamini−Hochberg corrected across the three comparisons within each panel. **Middle and Bottom rows**: Volcano plots and MA-plots for each CUT&Tag vs ChIP-seq comparison with each bin colored by its GC content, Z-scored DNase-seq or Z-scored ATAC-seq signal. Z-score signal for DNase-seq and ATAC-seq was calculated for each cell type independently.

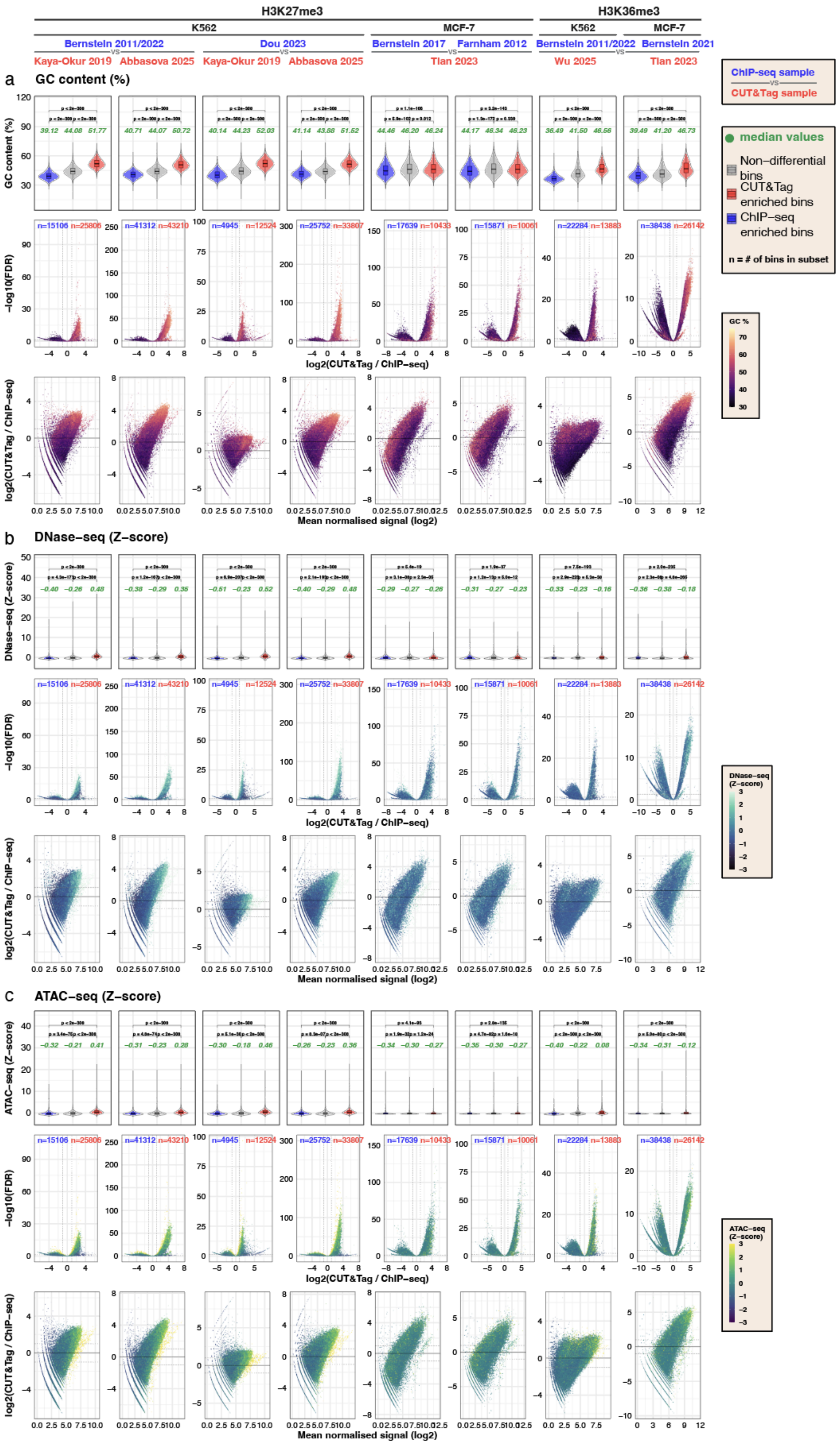

H3K27me3
H3K36me3
K562
MCF-7
K562
MCF-7
Bernstein 2011/2022
Dou 2023
Bernstein 2017
Farnham 2012
Bernstein 2011/2022
Bernstein 2021
vs
Kaya-Okur 2019
Abbasova 2025
Kaya-Okur 2019
Abbasova 2025
Tian 2023
Wu 2025
Tian 2023
ChIP-seq sample
vs
CUT&Tag sample
median values
Non-differential bins
CUT&Tag enriched bins
ChIP-seq enriched bins
n = # of bins in subset
a
GC content (%)
GC content (%)
-log10(FDR)
log2(CUT&Tag / ChIP-seq)
log2(CUT&Tag / ChIP-seq)
Mean normalised signal (log2)
GC %
b
DNase-seq (Z-score)
DNase-seq (Z-score)
DNase-seq (Z-score)
c
ATAC-seq (Z-score)
ATAC-seq (Z-score)
ATAC-seq (Z-score)

*(high resolution images attached separately as PDF)*

**Supplementary Figure 5.** ***Differential signals between CUT&Tag and ChIP-seq for broad marks H3K27me3 and H3K36me3 are associated with GC content and chromatin accessibility across multiple datasets in K562 and MCF-7 cell lines*.**

**a-c. Top row:** Violin plots showing the distribution of GC content (**a**) Z-scored DNase-seq signal (**b**) or Z-scored ATAC-seq signal (**c**) in ChIP-seq-enriched, CUT&Tag-enriched, and non-differential bin subsets. H3K36me3 comparisons use Active Gene Body bins while H3K27me3 comparisons use Polycomb-repressed bins. Additional metadata and general processing metrics for CUT&Tag (red) and ChIP-seq (blue) samples compared are available in **Supp Table 1,2**. Median value is shown in green. Differential bins are classified by DESeq2 (FDR < 0.05 and |log2FC| > 1.0). Adjusted P-value for the differences in distribution of GC content or Z-scored DNase-seq or Z-scored ATAC-seq signal between the three bin subsets is calculated with a two-sided Wilcoxon rank−sum test, Benjamini−Hochberg corrected across the three comparisons within each panel. **Middle and Bottom rows**: Volcano plots and MA-plots for each CUT&Tag vs ChIP-seq comparison with each bin colored by its GC content, Z-scored DNase-seq or Z-scored ATAC-seq signal. Z-scored signal for DNase-seq and ATAC-seq was calculated for each cell type and region set (e.g., K562 Active Gene Body bins) independently.

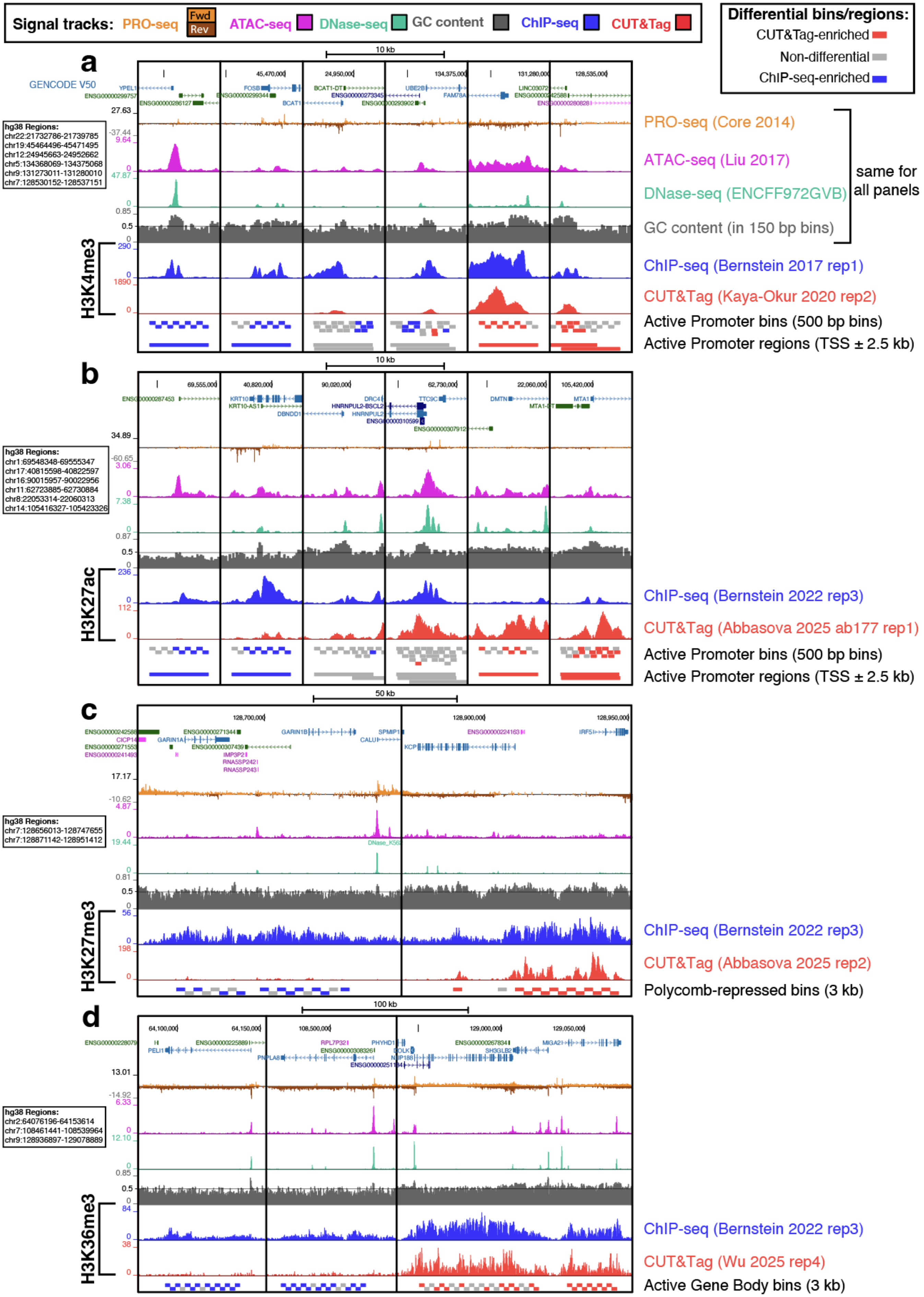
Signal tracks: PRO-seq Fwd Rev ATAC-seq DNase-seq GC content ChIP-seq CUT&Tag
Differential bins/regions:
CUT&Tag-enriched
Non-differential
ChIP-seq-enriched
a
10 kb
GENCODE V50
hg38 Regions:
chr22:21732786-21739785
chr19:45464496-45471495
chr12:24945663-24952662
chr5:134368069-134375068
chr9:131273011-131280010
chr7:128530152-128537151
PRO-seq (Core 2014)
ATAC-seq (Liu 2017)
DNase-seq (ENCFF972GVB)
GC content (in 150 bp bins)
same for all panels
H3K4me3
ChIP-seq (Bernstein 2017 rep1)
CUT&Tag (Kaya-Okur 2020 rep2)
Active Promoter bins (500 bp bins)
Active Promoter regions (TSS ± 2.5 kb)
b
10 kb
hg38 Regions:
chr1:69548348-69555347
chr17:40815598-40822597
chr16:90015957-90022956
chr11:62723885-62730884
chr8:22053314-22060313
chr14:105416327-105423326
H3K27ac
ChIP-seq (Bernstein 2022 rep3)
CUT&Tag (Abbasova 2025 ab177 rep1)
Active Promoter bins (500 bp bins)
Active Promoter regions (TSS ± 2.5 kb)
c
50 kb
hg38 Regions:
chr7:128656013-128747655
chr7:128871142-128951412
H3K27me3
ChIP-seq (Bernstein 2022 rep3)
CUT&Tag (Abbasova 2025 rep2)
Polycomb-repressed bins (3 kb)
d
100 kb
hg38 Regions:
chr2:64076196-64153614
chr7:108461441-108539964
chr9:128936897-129078889
H3K36me3
ChIP-seq (Bernstein 2022 rep3)
CUT&Tag (Wu 2025 rep4)
Active Gene Body bins (3 kb)

**Supplementary Figure 6.** ***Genome browser images showing loci of differential signal between ChIP-seq and CUT&Tag in K562 cells.*** ChIP-seq signal and enriched bins/regions are shown in blue while CUT&Tag signal and enriched bins/regions are shown in red; Non-differential bins/regions are in grey. PRO-seq is shown in Orange, ATAC-seq in magenta, DNase-seq in teal and average GC content in 150bp bins is shown in dark grey.

**a.** H3K4me3 ChIP-seq and CUT&Tag differential signals on Active Promoter bins (500bp) and full Active Promoter regions (TSS ±2.5 kb). Interactive session link: https://genome.ucsc.edu/s/emodolo/H3K4me3_K562_supp_fig_modolo_2026_multi%2Dregion

**b.** H3K27ac ChIP-seq and CUT&Tag differential signals on Active Promoter bins (500bp) and full Active Promoter regions (TSS ±2.5 kb). Interactive session link: https://genome.ucsc.edu/s/emodolo/H3K27ac_K562_Supp_fig_modolo_2026_multi%2Dregion

**c.** H3K27me3 ChIP-seq and CUT&Tag differential signals on Polycomb-repressed bins (3 kb). Interactive session link: https://genome.ucsc.edu/s/emodolo/H3K27me3_K562_Supp_fig_modolo_2026_multi%2Dregion

**d.** H3K36me3 ChIP-seq and CUT&Tag differential signals on Active Gene Body bins. Interactive session link: https://genome.ucsc.edu/s/emodolo/H3K36me3_K562_Supp_fig_modolo_2026_multi%2Dregion

**Supp. Data 1-4** provide coordinates for inspection of additional differential analysis results using the interactive browser sessions.

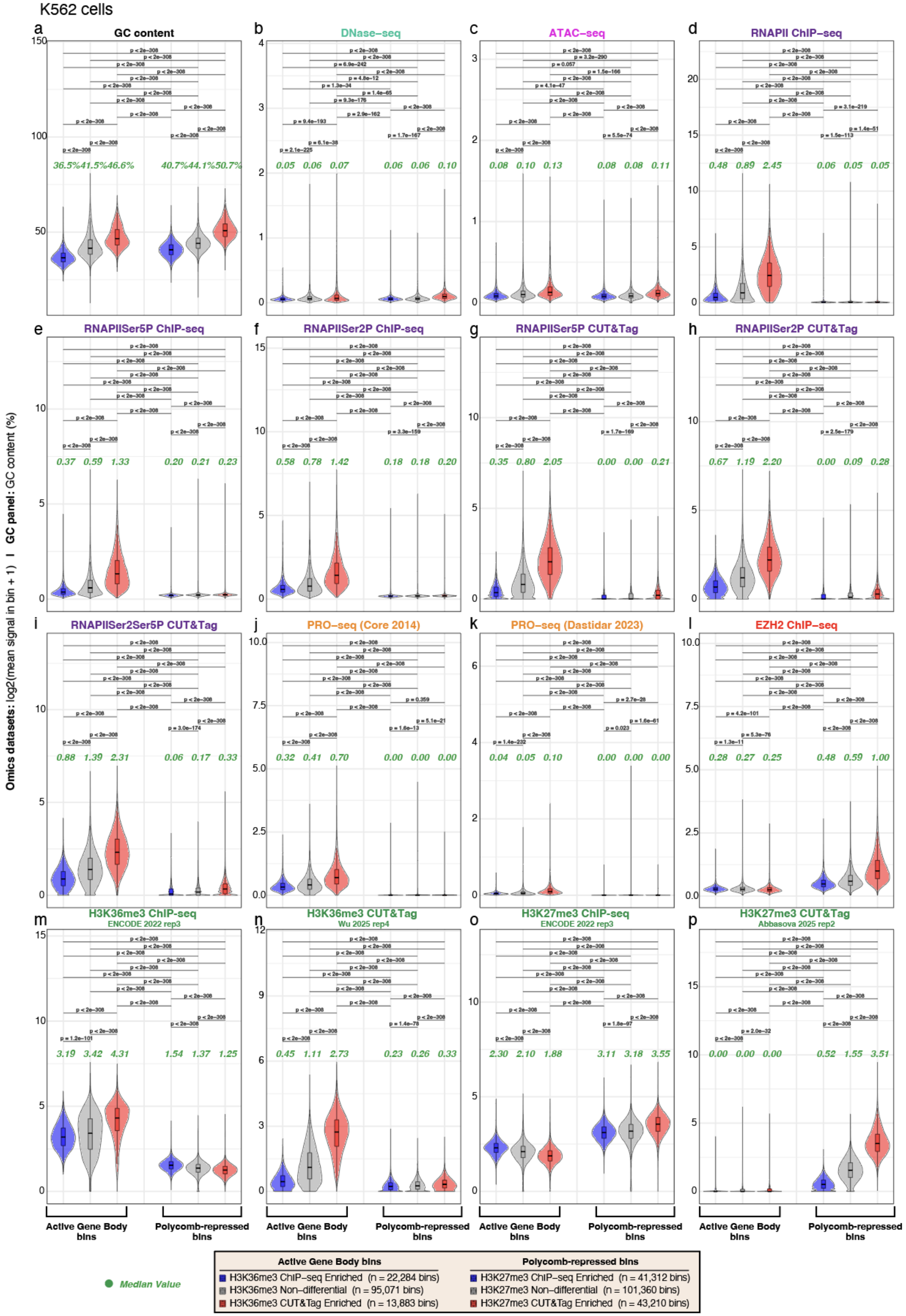

K562 cells
a GC content
b DNase–seq
c ATAC–seq
d RNAPII ChIP–seq
e RNAPIISer5P ChIP-seq
f RNAPIISer2P ChIP-seq
g RNAPIISer5P CUT&Tag
h RNAPIISer2P CUT&Tag
i RNAPIISer2Ser5P CUT&Tag
j PRO–seq (Core 2014)
k PRO–seq (Dastidar 2023)
l EZH2 ChIP–seq
m H3K36me3 ChIP–seq ENCODE 2022 rep3
n H3K36me3 CUT&Tag Wu 2025 rep4
o H3K27me3 ChIP–seq ENCODE 2022 rep3
p H3K27me3 CUT&Tag Abbasova 2025 rep2
Omics datasets: log2(mean signal in bin + 1) | GC panel: GC content (%)
Active Gene Body bins
Polycomb-repressed bins
Median Value
Active Gene Body bins
H3K36me3 ChIP–seq Enriched (n = 22,284 bins)
H3K36me3 Non–differential (n = 95,071 bins)
H3K36me3 CUT&Tag Enriched (n = 13,883 bins)
Polycomb-repressed bins
H3K27me3 ChIP–seq Enriched (n = 41,312 bins)
H3K27me3 Non–differential (n = 101,360 bins)
H3K27me3 CUT&Tag Enriched (n = 43,210 bins)

*(high resolution images attached separately as PDF)*

**Supplementary Figure 7. *Distribution of reference dataset as well as ChIP-seq and CUT&Tag signal per-bin across differentially enriched Active Gene Body and Polycomb-repressed bins in K562 cells.***

***a-p.*** Left block: active gene-body bins (3 kb) classified by the H3K36me3 CUT&Tag vs ChIP-seq differential comparison. Right block: Polycomb-repressed bins (3 kb) classified by the equivalent H3K27me3 comparison. Within each block, violin plots represent the distribution of signal on bins that are ChIP-seq-enriched (blue), non-differential (grey), or CUT&Tag-enriched (red). Each of the 16 panels is one dataset plotted over all six bin subsets. Violins show mean signal per bin, log2(x + 1) transformed for each omics dataset, while the GC panel shows mean GC content as a percentage, untransformed. Violins are scaled to equal width; the inset box plot marks the interquartile range and median, and the green italic number states the median signal of the bin subset. The legend shows the number of bins per bin subset. Brackets carry Benjamini–Hochberg-adjusted p-values from two-sided Wilcoxon rank-sum tests over all fifteen pairs of bin subsets, corrected within each panel.

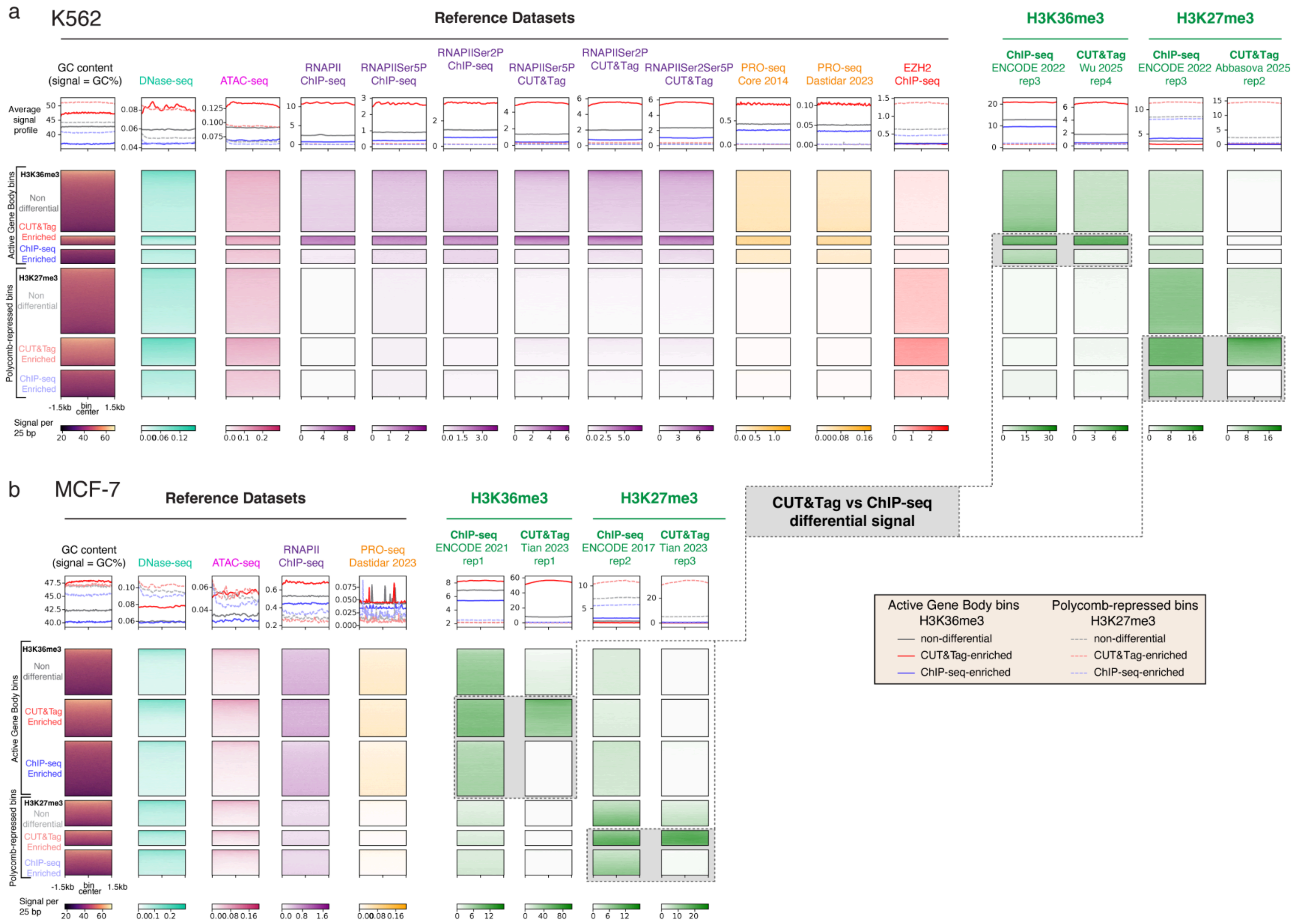


**Supplementary Figure 8. Regions showing limited H3K27me3 and H3K36me3 CUT&Tag signals exhibit low GC content and reduced chromatin accessibility in K562 and MCF-7 cells.**

Heatmaps on active gene body and Polycomb-repressed bin subsets generated by H3K36me3 and H3K27me3 differential analysis, respectively. Datasets are from K562 cells (**a**) and MCF-7 cells (**b**); heatmaps are computed by taking average signal in 25 bp bins across each 3 kb Active Gene Body or Polycomb-repressed bin, with the color ceiling set to each column's 97th percentile to suppress outliers. Each bin subset is individually sorted by decreasing GC content, and is centred on the 3 kb bin (1.5 kb flanks). Above each heatmap column, a line plots represent the average signal (or in the case of GC content, the GC%) in 25 bp bins across the 3 kb bins for each subset. Reference datasets for K562 cells include: GC content, DNase-seq, ATAC-seq, ChIP-seq and CUT&Tag for RNAPII(RNAPIISer2PSer5P), RNAPIISer5P and RNAPIISer2P, as well as PRO-seq from two sources[1,2], and ChIP-seq for EZH2. Reference datasets for MCF-7 cells include GC content, DNase-seq, ATAC-seq, RNAPII ChIP-seq, and PRO-seq[2]. Grey box: ChIP-seq and CUT&Tag signal (quantified from a representative sample) on differentially-enriched bins, classified by DESeq2 as having an FDR<0.05 and |Log2(CUT&Tag/ChIP-seq)| > 1.0 (**Methods**).

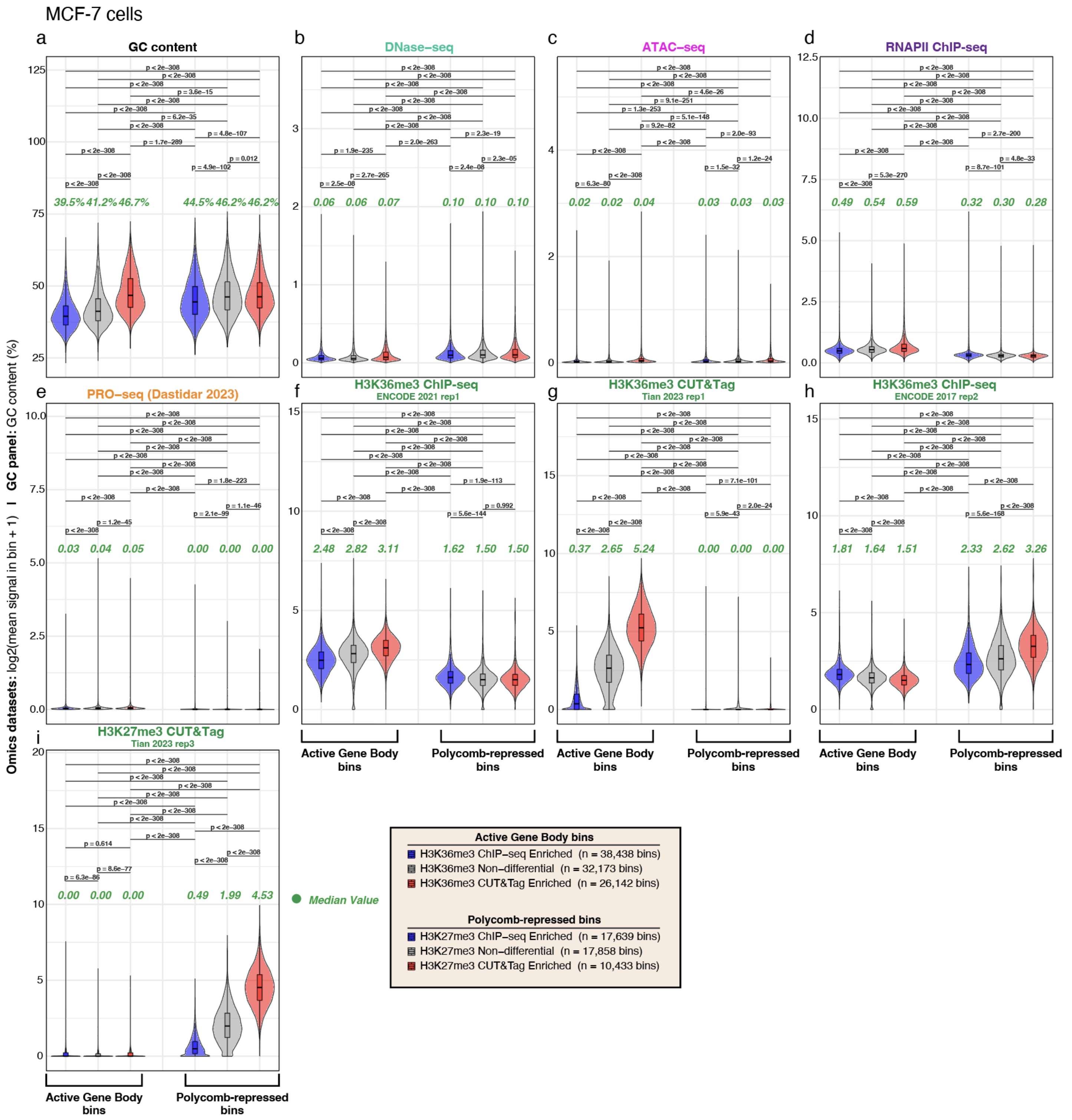


**Supplementary Figure 9.** ***Distribution of reference dataset as well as ChIP-seq and CUT&Tag signal per-bin across differentially enriched Active Gene Body and Polycomb-repressed bins in MCF-7 cells.***

***a-p.*** Left block: active gene-body bins (3 kb) classified by the H3K36me3 CUT&Tag vs ChIP-seq differential comparison. Right block: Polycomb-repressed bins (3 kb) classified by the equivalent H3K27me3 comparison. Within each block, violin plots represent the distribution of signal on bins that are ChIP-seq-enriched (blue), non-differential (grey), or CUT&Tag-enriched (red). Each of the 16 panels is one dataset plotted over all six bin subsets. Violins show mean signal per bin, log2(x + 1) transformed for each omics dataset, while the GC panel shows mean GC content as a percentage, untransformed. Violins are scaled to equal width; the inset box plot marks the interquartile range and median, and the green italic number states the median signal of the bin subset. The legend shows the number of bins per bin subset. Brackets carry Benjamini–Hochberg-adjusted p-values from two-sided Wilcoxon rank-sum tests over all fifteen pairs of bin subsets, corrected within each panel.

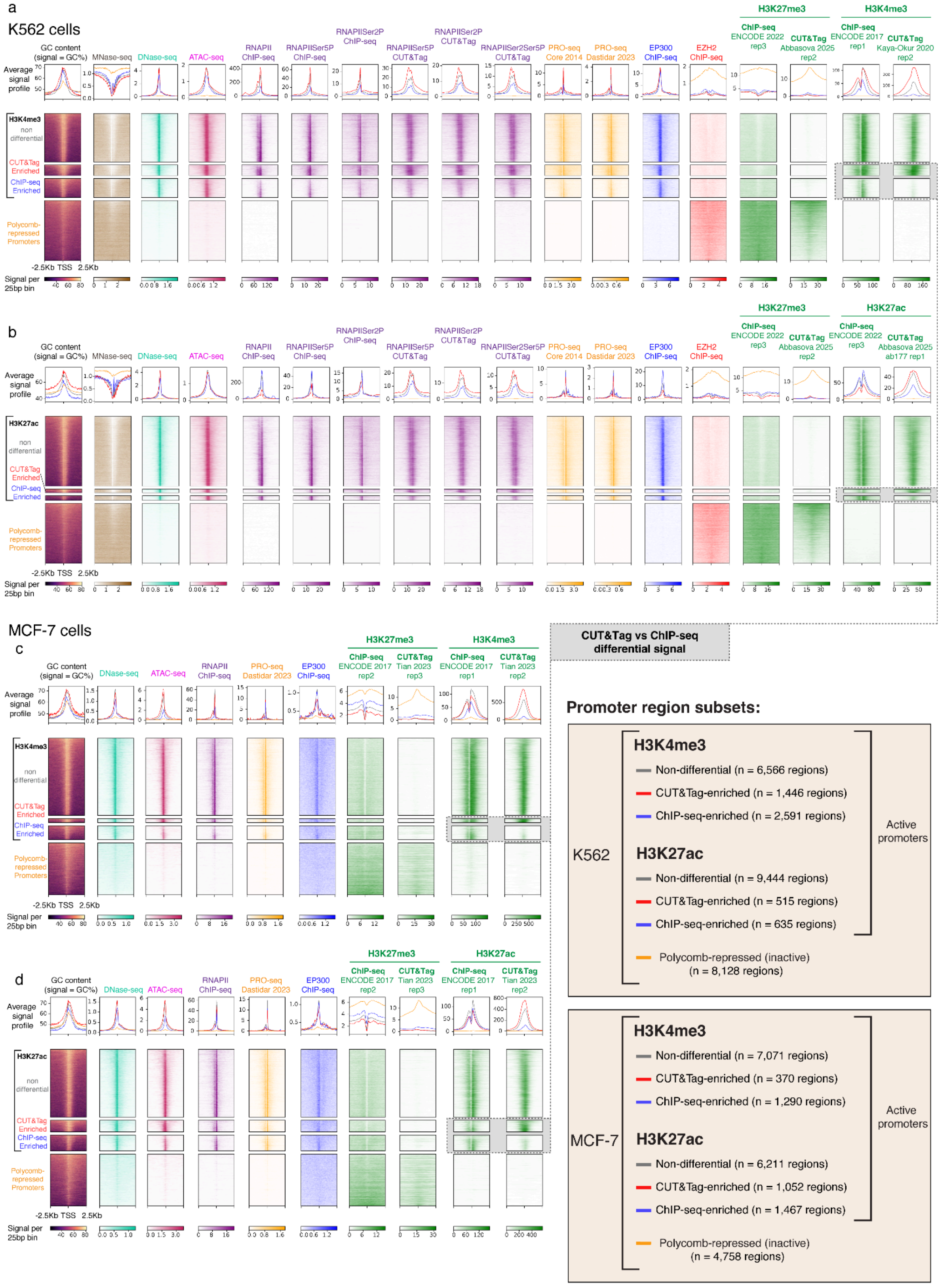

a
K562 cells
b
MCF-7 cells
c
d
CUT&Tag vs ChIP-seq differential signal
Promoter region subsets:
K562
H3K4me3
Non-differential (n = 6,566 regions)
CUT&Tag-enriched (n = 1,446 regions)
ChIP-seq-enriched (n = 2,591 regions)
H3K27ac
Non-differential (n = 9,444 regions)
CUT&Tag-enriched (n = 515 regions)
ChIP-seq-enriched (n = 635 regions)
Active promoters
Polycomb-repressed (inactive) (n = 8,128 regions)
MCF-7
H3K4me3
Non-differential (n = 7,071 regions)
CUT&Tag-enriched (n = 370 regions)
ChIP-seq-enriched (n = 1,290 regions)
H3K27ac
Non-differential (n = 6,211 regions)
CUT&Tag-enriched (n = 1,052 regions)
ChIP-seq-enriched (n = 1,467 regions)
Active promoters
Polycomb-repressed (inactive) (n = 4,758 regions)

*(high resolution images attached separately as PDF)*

**Supplementary Figure 10.** ***Active promoters showing differential H3K4me3 and H3K27ac ChIP-seq and CUT&Tag signals are associated with GC content and chromatin accessibility patterns in K562 and MCF-7 cells.***

For H3K4me3 and H3K27ac a differential analysis between ChIP-seq and CUT&Tag was performed on Active Promoter regions (TSS ±2.5 kb). Heatmaps show signals quantified on ChIP-seq-enriched, CUT&Tag-enriched and non-differential Active Promoter regions as well as Polycomb-repressed Promoters Regions (used as a set of background regions representing silenced promoters). H3K4me3 and H3K27ac differentially-enriched promoter regions classified by DESeq2 as having an FDR<0.05 and |Log2(CUT&Tag/ChIP-seq)| > 1.0. (**Methods**). **a,b** shows data from K562 cells, **c,d** shows data from MCF-7 cells; heatmaps are computed by taking average signal (or in the case of GC content, the GC%) in 25 bp bins across each Active or Polycomb-repressed Promoter Region (5kb), with color ceiling set to each column's 97th percentile to suppress outliers. Each Active Promoter region subset is individually sorted by decreasing GC content and centered on the TSS with 2.5 kb flanks. Above each heatmap column, Average signal profile (metagene) plots represent the average signal in 25 bp bins across the 5 kb region for each promoter region subset. Grey box: ChIP-seq and CUT&Tag signal on differentially-enriched Active Promoter regions.

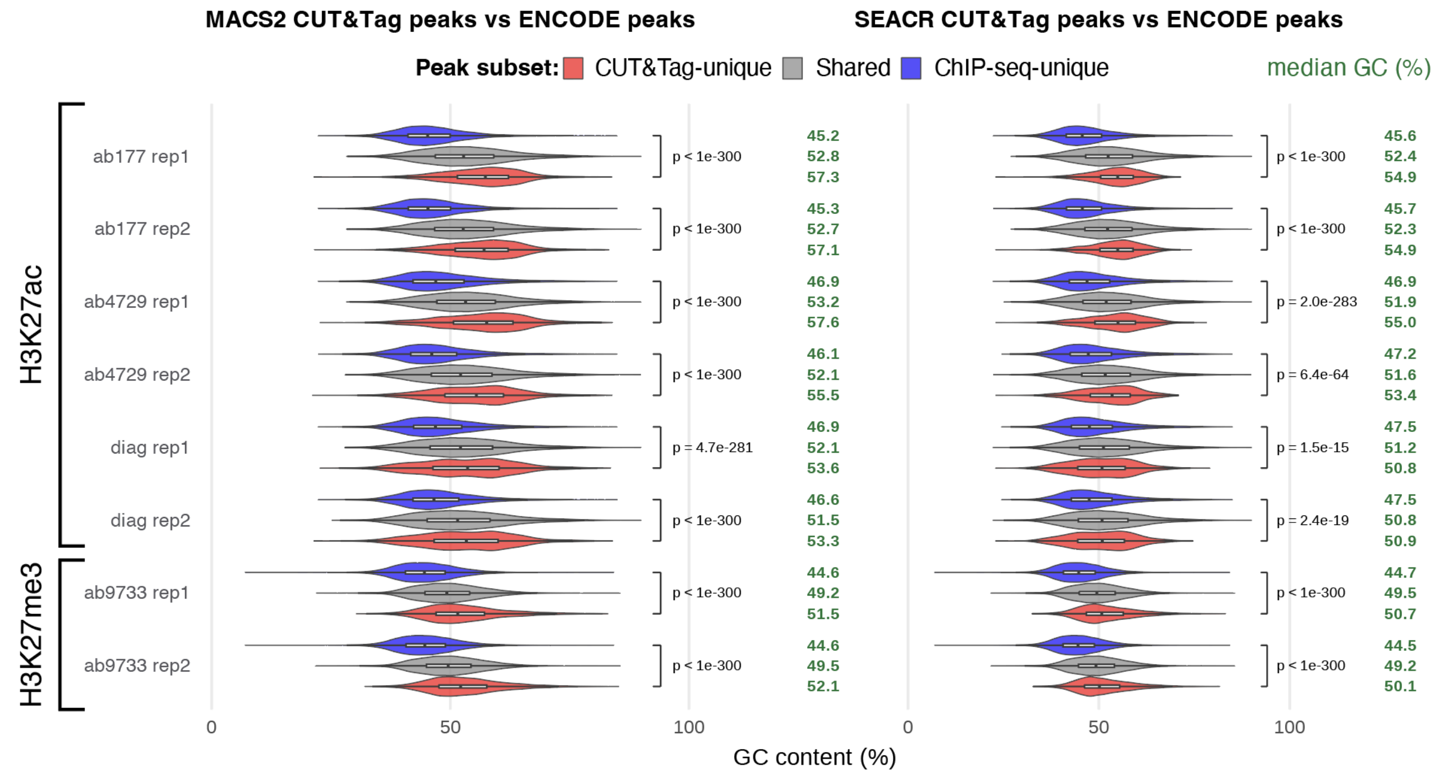


**Supplementary Figure 11.** ***ENCODE unique peaks are associated with lower GC content compared to CUT&Tag unique peaks for H3K27ac and H3K27me3.***

**a.** Peak overlap analysis comparing H3K27ac and H3K27me3 CUT&Tag peaks called by MACS2 and SEACR (generated by the Marzi group[3]) to ENCODE ChIP-seq peaks (*H3K27ac: ENCFF044JNJ, H3K27me3: ENCFF001SZF*), hg19 peaks used throughout. Antibody code and biological replicate number are shown for each CUT&Tag sample. Left panel: CUT&Tag peaks called with MACS2. Right panel: CUT&Tag peaks called with SEACR. Violins show the distribution of per-peak GC content (i.e., mean GC% per peak) and are scaled to equal width; the inset white box plot marks the interquartile range and median. Green values on the right of each violin state the median GC content of each peak subset. Brackets carry the two-sided Wilcoxon rank-sum p-value for the CUT&Tag-unique versus ChIP-seq-unique comparison, Benjamini–Hochberg-adjusted within each panel.

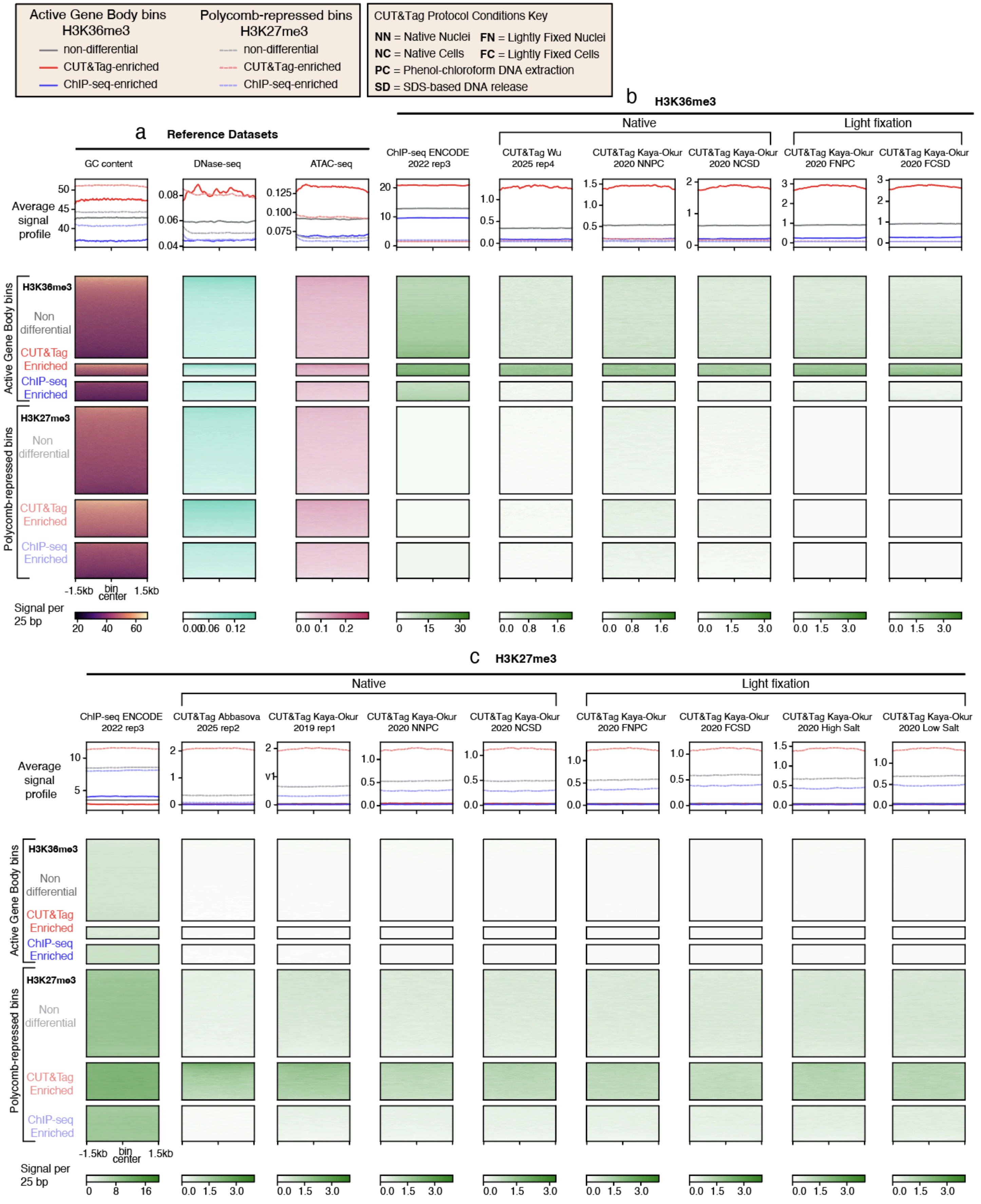
Active Gene Body bins H3K36me3
non-differential
CUT&Tag-enriched
ChIP-seq-enriched
Polycomb-repressed bins H3K27me3
non-differential
CUT&Tag-enriched
ChIP-seq-enriched
CUT&Tag Protocol Conditions Key
NN = Native Nuclei
FN = Lightly Fixed Nuclei
NC = Native Cells
FC = Lightly Fixed Cells
PC = Phenol-chloroform DNA extraction
SD = SDS-based DNA release
a Reference Datasets
GC content
DNase-seq
ATAC-seq
Average signal profile
b H3K36me3
Native
Light fixation
ChIP-seq ENCODE 2022 rep3
CUT&Tag Wu 2025 rep4
CUT&Tag Kaya-Okur 2020 NNPC
CUT&Tag Kaya-Okur 2020 NCSD
CUT&Tag Kaya-Okur 2020 FNPC
CUT&Tag Kaya-Okur 2020 FCSD
Active Gene Body bins
H3K36me3
Non differential
CUT&Tag Enriched
ChIP-seq Enriched
Polycomb-repressed bins
H3K27me3
Non differential
CUT&Tag Enriched
ChIP-seq Enriched
-1.5kb
bin center
1.5kb
Signal per 25 bp
c H3K27me3
Native
Light fixation
ChIP-seq ENCODE 2022 rep3
CUT&Tag Abbasova 2025 rep2
CUT&Tag Kaya-Okur 2019 rep1
CUT&Tag Kaya-Okur 2020 NNPC
CUT&Tag Kaya-Okur 2020 NCSD
CUT&Tag Kaya-Okur 2020 FNPC
CUT&Tag Kaya-Okur 2020 FCSD
CUT&Tag Kaya-Okur 2020 High Salt
CUT&Tag Kaya-Okur 2020 Low Salt
Average signal profile
Signal per 25 bp

**Supplementary Figure 12. CUT&Tag for H3K27me3 and H3K36me3 in K562 cells shows limited sensitivity in low GC regions compared to ChIP-seq across a variety of CUT&Tag protocol parameters.**

H3K36me3 and H3K27me3 CUT&Tag samples prepared using various starting sample and fixation conditions (NN = Native Nuclei, FN = lightly Fixed Nuclei, NC = Native Cells, FC = lightly Fixed Cells) as well as DNA isolation and library prep methods (PC = Phenol-chloroform DNA extraction and PCR in new tube, SD = SDS-based release of DNA fragments, PCR performed in the same tube as previous steps), and conditions during tagmentation steps (Low Salt = 10mM TAPS, High Salt = 300mM NaCl) were quantified on previously identified differential regions (more details on protocol variations in Kaya-Okur et al. 2020[4]). Differential (i.e., CUT&Tag-enriched, ChIP-seq-enriched) and non-differential bin subsets for Active Gene Body and Polycomb-repressed bins were identified by performing a differential analysis with DESeq2 comparing ChIP-seq and CUT&Tag signal for H3K36me3 and H3K27me3 on each bin set respectively (same differential regions identified in **Fig. 1d,e** and displayed in **Fig. 2** and **Supp. Fig. 8**).

**a.** Quantification of reference dataset signal including GC content, DNase-seq and ATAC-seq.

**b-c)** Quantification of ChIP-seq and CUT&Tag samples for H3K36me3 (**b**) and H3K27me3 (**c**). Heatmaps are computed by taking average signal in 25 bp bins across each 3 kb Active Gene Body or Polycomb-repressed bin, with the color ceiling set to each column's 97th percentile to suppress outliers. Each bin subset is individually sorted by decreasing GC content, and is centred on the 3 kb bin (1.5 kb flanks). Above each heatmap column, a line plots represent the average signal (or in the case of GC content, the GC%) in 25 bp bins across the 3 kb bins for each subset. All CUT&Tag samples were downsampled to 1 million fragments for fair comparison across CUT&Tag protocol parameters.
