## Supplementary figures and images for "Discrepancies between ChIP-seq and CUT&Tag histone mark profiles are explained by GC content and chromatin accessibility"

### Supplemental Figure 3 (high resolution)

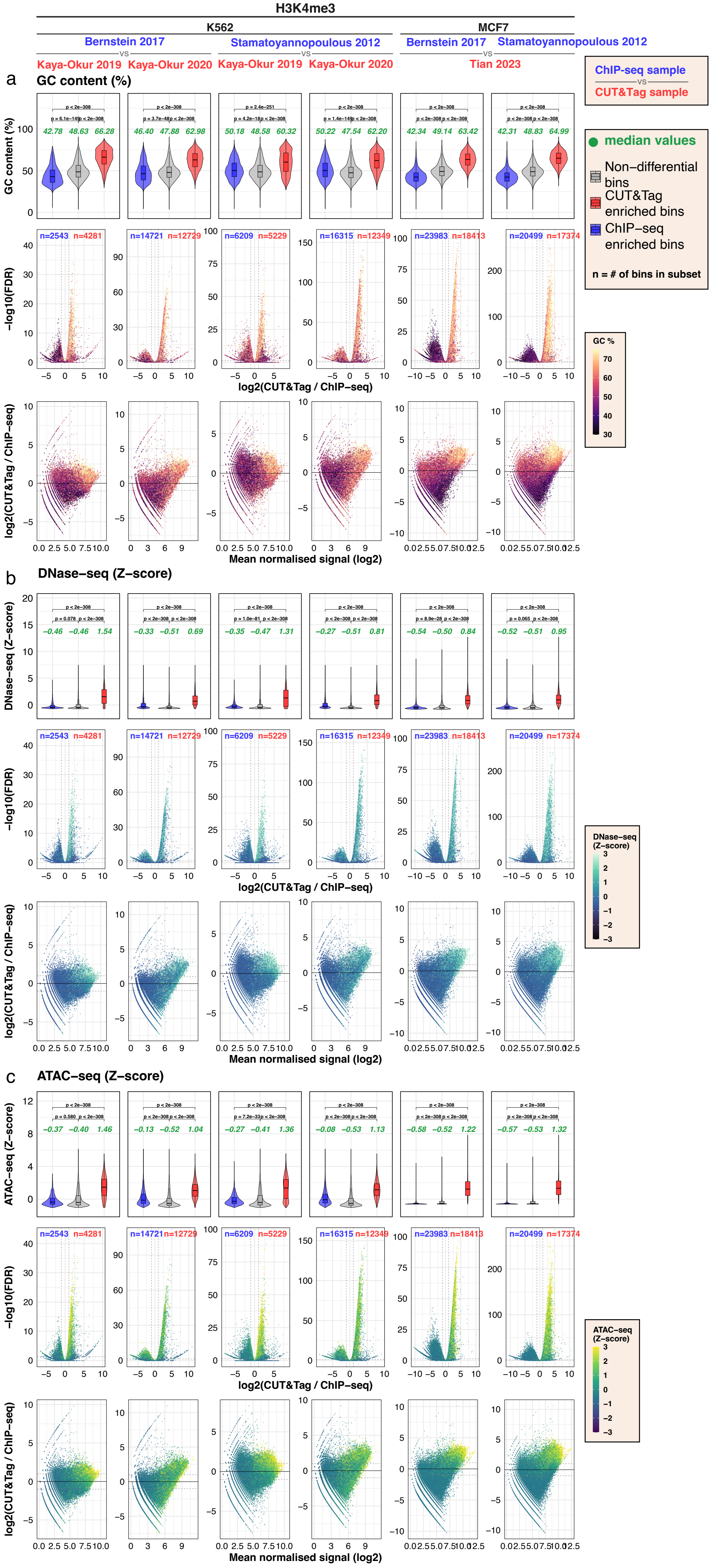

### Supplemental Figure 4 (high resolution)

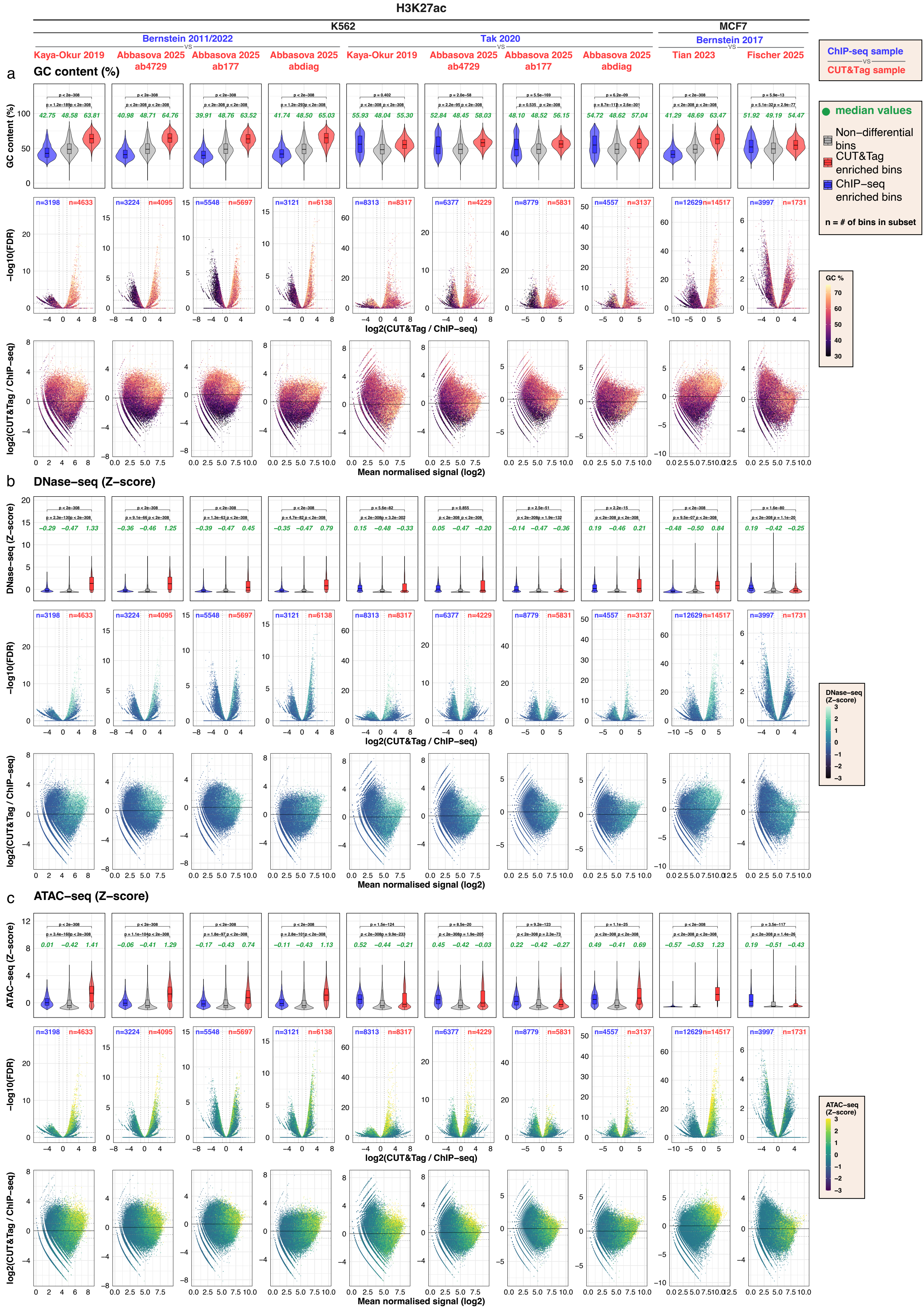

### Supplemental Figure 5 (high resolution)

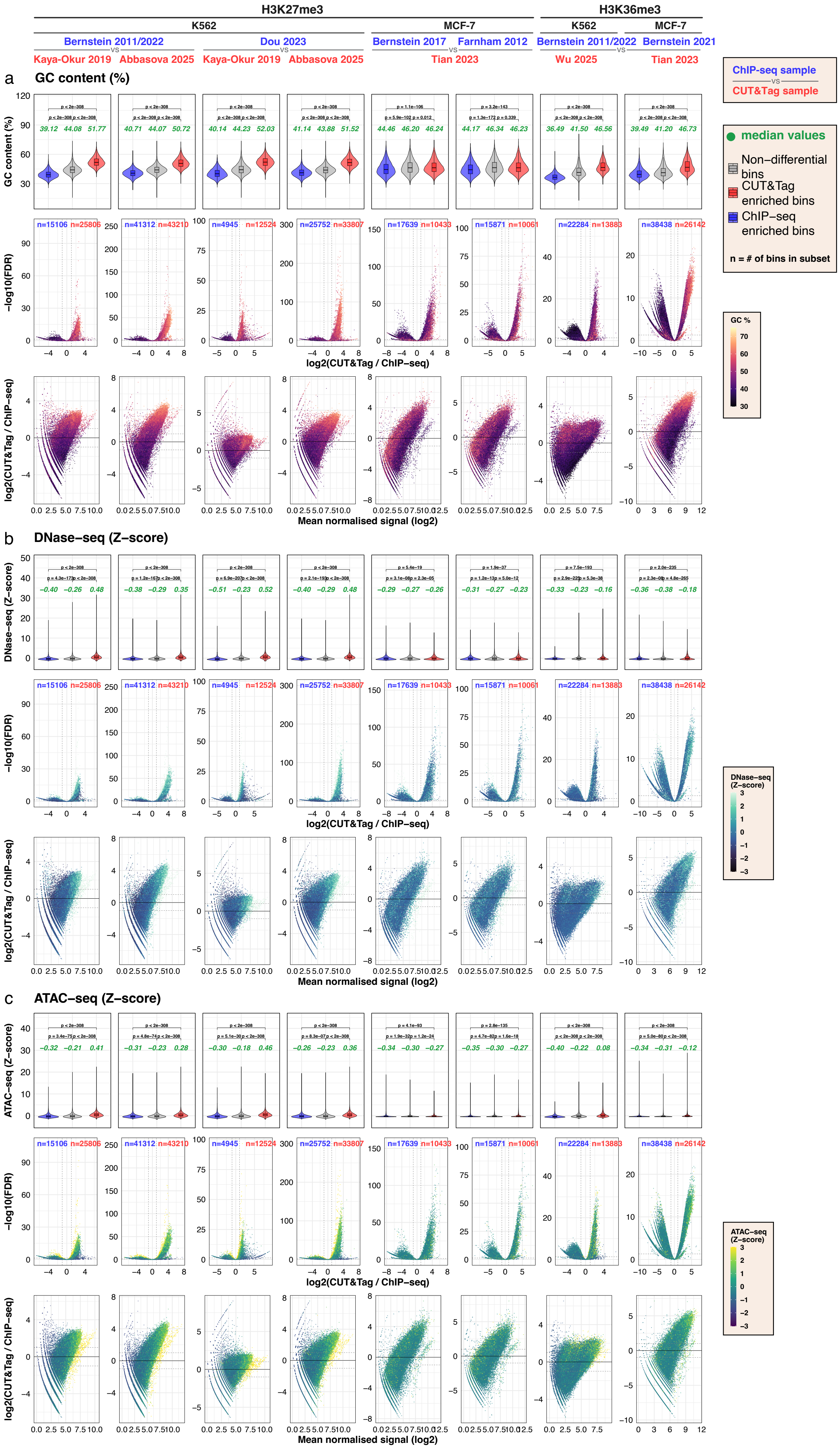

### Supplemental Figure 7 (high resolution)

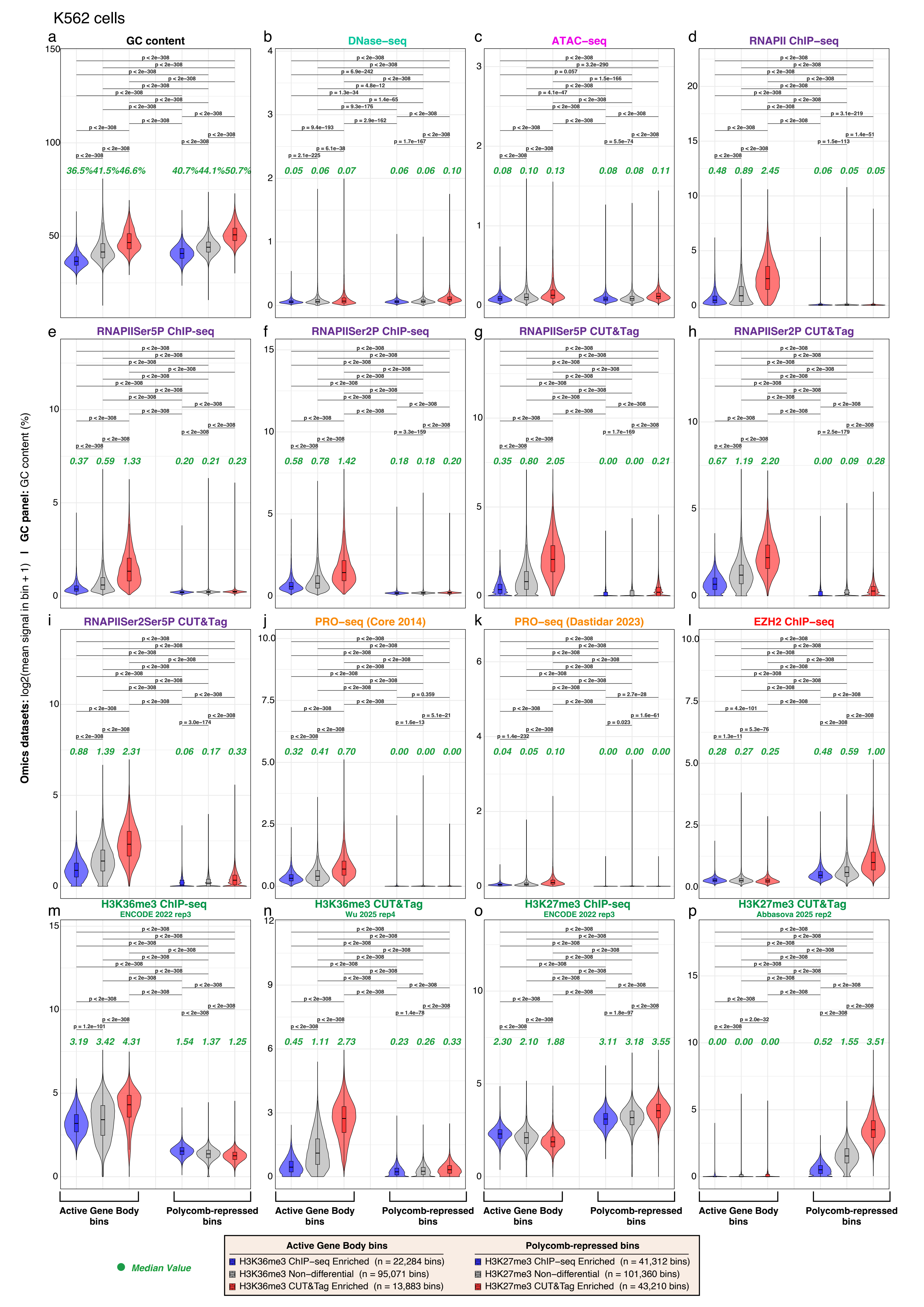

### Supplemental Figure 10 (high resolution)

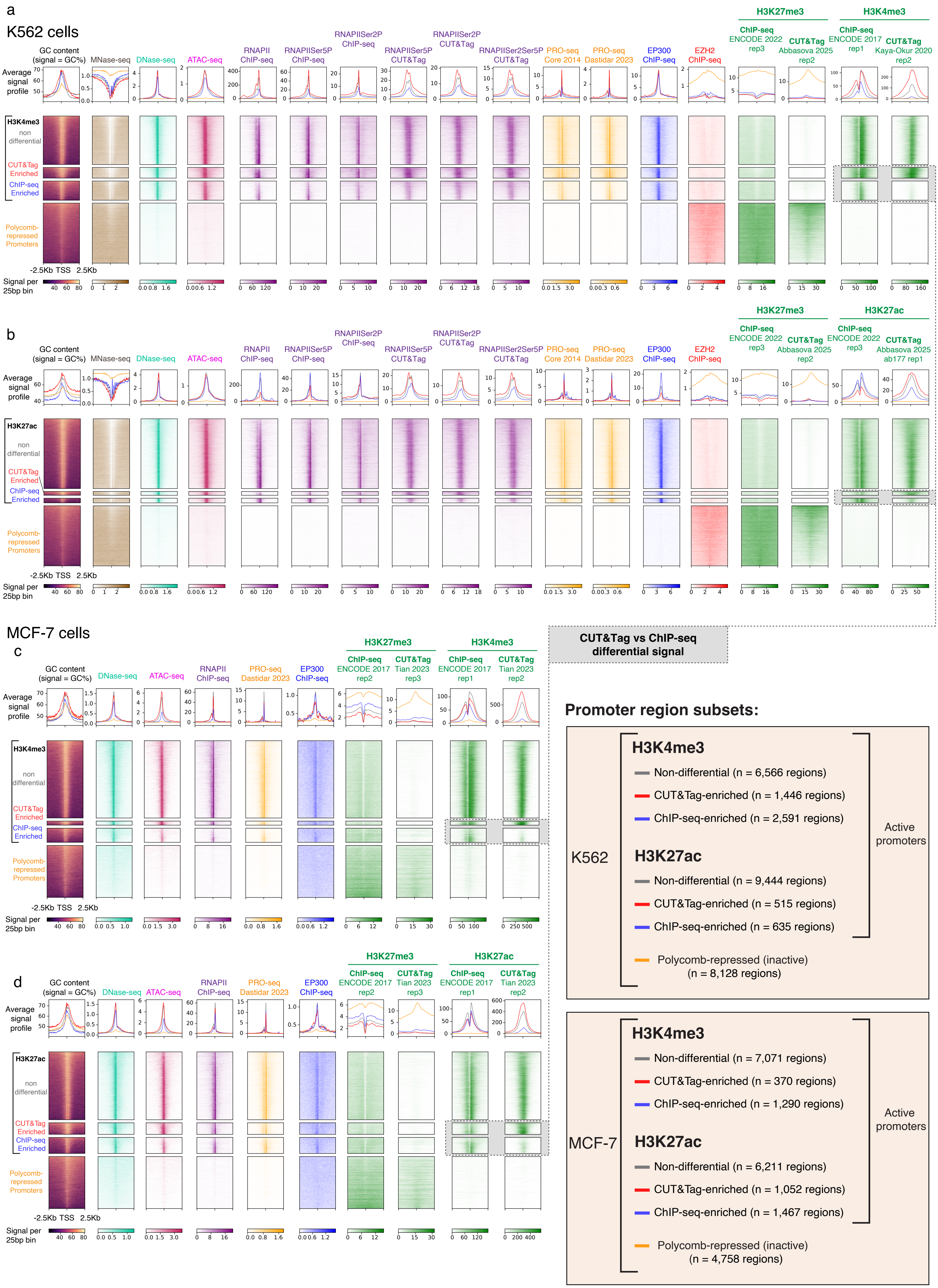
